# Mechanistic insights into *E. coli* recovery from growth arrest

**DOI:** 10.64898/2026.01.09.698670

**Authors:** Ahmed Hassan, Yuko Nakano, Howard Gamper, Isao Masuda, Matyas Pinkas, Sathya Nagarajan, Jonathan Dworkin, Gregor Blaha, Ya-Ming Hou, Gabriel Demo

## Abstract

Bacteria survive hostile conditions in clinically relevant conditions by shutting down protein synthesis, but how they restart growth remains poorly understood. Here, we use an *E. coli* Δ*rimM* strain, which exhibits a prolonged growth arrest, as a model to investigate how bacteria recover from this arrested state and restore protein synthesis. RimM is a conserved ribosome maturation factor for the 3’-major (head) domain of the 16S rRNA within the bacterial 30S subunit. The loss of RimM causes a significantly longer delay in recovery than other 30S maturation factors, including RbfA – the presumed primary factor in 30S maturation. Cryo-EM analysis of Δ*rimM* ribosomes revealed a delayed recruitment of ribosomal proteins to the 30S head domain and increased occupancy of the initiation factors IF1 and IF3, as well as recruitment of the silencing factor RsfS to the 50S subunit. These coordinated changes provide a safeguarding mechanism to block the assembly of premature 70S ribosomes. Notably, while the delayed 30S assembly in Δ*rimM* reduces the activity of global protein synthesis during the recovery phase, bacteria attempt to compensate for this deficiency by producing higher levels of the ribosomal machinery, indicating a programmatic change in energy allocation to generate the ribosome machinery. These findings highlight the importance of the RimM-assisted assembly of the ribosomal head domain for bacterial recovery from growth arrest.

## Introduction

Bacteria arrest growth under stress conditions, many of which are relevant to clinical settings^1^. In growth arrest, bacteria terminate ribosome-mediated protein synthesis and dissociate ribosomes from the associated mRNA and tRNAs. Although bacteria have evolved a wide range of strategies to exit growth arrest^2,3^, one of the key elements is to re-assemble the ribosome to resume protein synthesis. However, the mechanisms by which bacterial ribosomes recover from growth arrest remain poorly understood, creating a critical gap in our knowledge of stress adaptation and posing a major challenge to develop effective treatments of bacterial infections. To address this gap, we use an *E. coli* Δ*rimM* strain, which exhibits a prolonged growth arrest^4,5^, as a model to identify factors that determine ribosome recovery.

RimM^6^ is an evolutionarily conserved ribosomal maturation factor for assembly of the bacterial 30S small ribosomal subunit. The assembly of the 30S begins with transcriptional synthesis of the 17S precursor of the 16S rRNA, which is then processed, post-transcriptionally modified, and folded into distinct structural domains^7^. The folding of the 16S rRNA occurs in coordination with maturation factors to guide the sequential recruitment^8,9^ of ribosomal proteins (r-proteins) to the 30S subunit. This process produces a series of pre-30S intermediates^8–10^ *en route* to the fully mature state of the 30S subunit. Immature assembly of the 30S would produce error-prone ribosomes that disrupt the homeostasis of protein synthesis^11,12^. Based on the known maturation process of the 30S subunit^9,10^, ribosome binding factor A (RbfA) is considered to be the main organizer among the maturation factors of the 30S subunit (Fig. 1a). It is one of the first to engage in the 30S maturation pathway and it remains involved until the completion of maturation^13,14^, being released by the late-acting assembly factor RsgA^15^ and displaced by initiation factor IF3^14,16^. RbfA is required for two important stages of maturation – formation of the central pseudoknot structure of the 16S rRNA in the early stage^13^, and the docking of helix 44 into the decoding site in a later stage^17^. In contrast, RimM comes after RbfA in the maturation process and has a shorter residence time on pre-30S particles^18^. It is specifically involved in proper folding of the 3’-major domain of the 16S rRNA localized in the 30S head domain^5,13,19^, where it recruits secondary- and tertiary-level r-proteins to stabilize assembly intermediates^4,5,13,20^.

**Figure 1.**
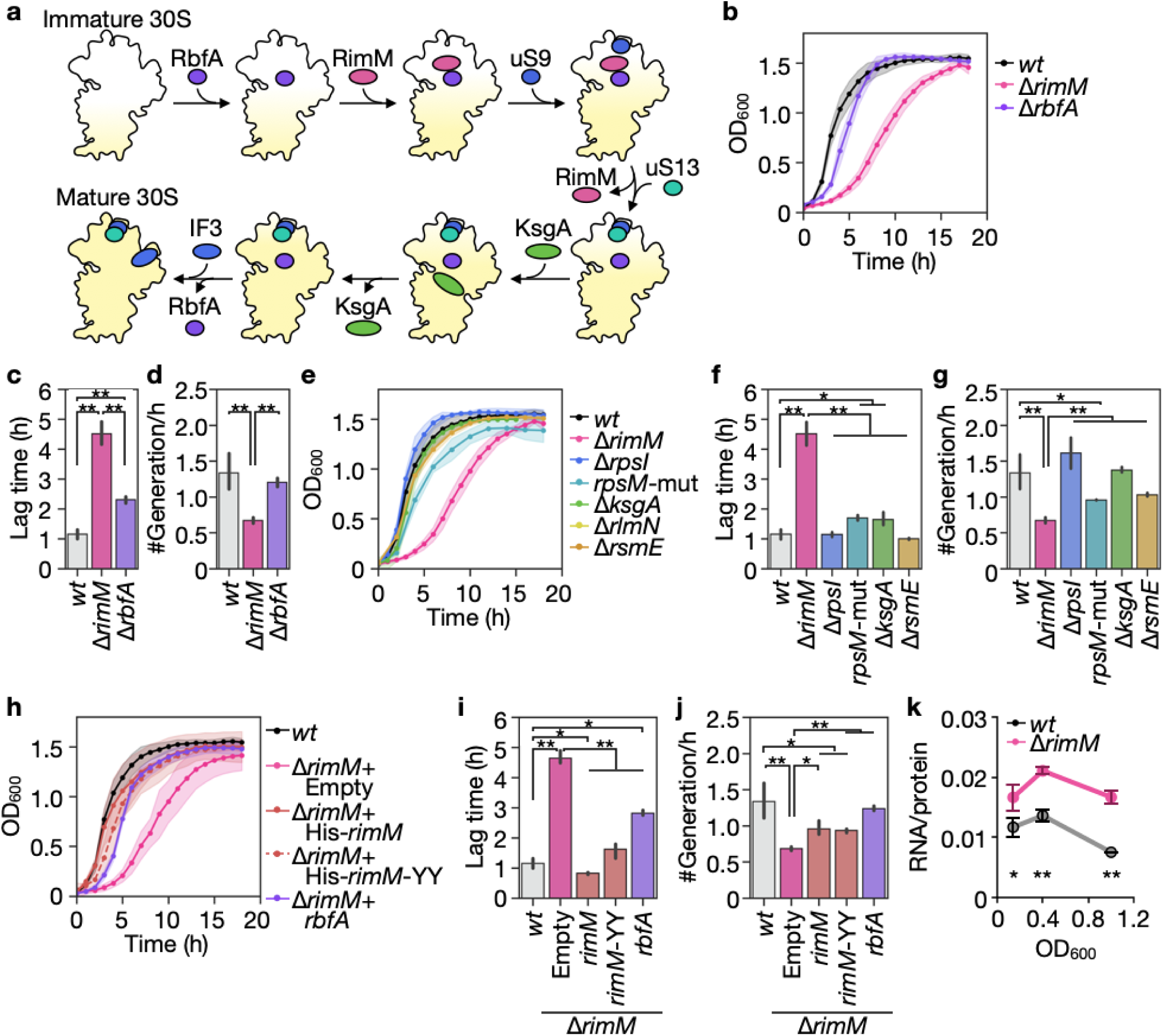
Assembly of *E. coli* 30S subunit in growth recovery. **(a)** Scheme of the 30S subunit assembly highlighting the biogenesis factors and r-proteins studied in this study. These include RbfA, RimM, uS9, uS13, KsgA, and IF3. The entry and exit points of RsmE in the pathway are not shown, due to lack of information. **(b)** Growth profiles of the *wt* strain MG1655 (n = 13), Δ*rimM* (n = 13), and Δ*rbfA* (n =3) in LB at 37°C after 1 day pre-culture. **(c)** Lag time calculated from OD600 of cultures in (b), as the time required to reach OD600 of 0.2. **(d)** Growth rates (the number of generations per hour) of cultures in (b). Error bars, mean ± 95% confidence interval. \**p* < 0.05, \*\**p* < 0.01; two-tailed Welch’s *t*-test. **(e)** Growth profiles of *wt* strain MG1655 (n = 13), Δ*rimM* (n = 13), Δ*rpsI* (lacking uS9) (n = 6), *rpsM*-mut (an uS13 mutant, lacking residues 82-117) (n = 3), Δ*ksgA* (n = 3), and Δ*rsmE* (n = 3) in LB at 37°C after 1 day of pre-culture. **(f)** Lag time calculated from OD600 of cultures in (e), as the time required to reach OD600 of 0.2. **(g)** Growth rates (the number of generations per hour) of cultures in (e). **(h)** Growth profiles of *wt* and Δ*rimM* MG1655 strains in LB at 37°C after 1 day of pre-culture. The gene for His-*rimM* (n = 3), His-*rimM*-*YY* mutant (His-*rimM*-Y106A/Y107A) (n = 3), or *rbfA* (n = 3) was overexpressed in Δ*rimM*, and an empty vector was used as a negative control. **(i)** Lag time calculated from OD600 of cultures in (h), as the time required to reach OD600 of 0.2. **(j)** Growth rates (the number of generations per hour) of cultures in (h). **(k)** Ribosome contents of the MG1655 *wt* (n = 4) and Δ*rimM* strains (n = 3). \**p* < 0.05, \*\**p* < 0.01; two-tailed Student’s *t*-test.

Notably, RimM is not essential for bacterial survival, as several r-proteins (e.g., uS2, uS3, uS10, uS13, uS14, and uS19) can spontaneously bind to and facilitate the folding of the head region of the 16S rRNA^4,5^. However, *E. coli* strains lacking Δ*rimM* exhibited a markedly slow growth phenotype in previous studies^4,5^, although no direct comparisons of the growth recovery of Δ*rimM* with Δ*rbfA* have been reported, nor with *E. coli* strains lacking other 30S maturation factors and r-proteins. It is also unknown whether Δ*rimM* or the deletion of any 30S assembly factors induces coordinated changes in the maturation process of the large ribosomal subunit 50S, which also occurs through the stepwise incorporation of r-proteins into the 23S and 5S rRNA intermediates to form functional domains of the peptidyl transferase center and the central protuberance^21–24^. Late-stage 50S maturation involves stable interactions of the 23S rRNA with r-proteins, the assembly of a 5S complex with r-proteins, and the formation of inter-subunit bridge precursors^21,23^. Final maturation of the 50S subunit includes conformational rearrangements and post-transcriptional modifications of both 23S and 5S rRNAs, which together generate an active 50S subunit^23,24^. It is noteworthy that ribosomal silencing factor S (RsfS)^25,26^, which coordinates with other late-stage biogenesis factors (e.g., RluD, YjgA, and ObgE)^25^ stabilize essential regions of the 23S rRNA and recruit r-proteins to complete 50S assembly. Under stress conditions^26,27^, RsfS interacts with r-protein uL14 at the subunit interface to block immature pre-50S particles from joining a 30S subunit^25^. While the conservation of RsfS across bacterial species^25–27^ highlights its role in 50S assembly and function^3,28^, how it responds to the accumulation of pre-30S particles is unknown. Although bacterial ribosomal structures lacking specific maturation factors^4,13^, including pre-30S structures lacking RimM^5^, are available, to date there is no complete set of *ex vivo* structures of 30S, 50S, and 70S particles from a Δ*rimM* strain. This limits our understanding of how these particles coordinate to assemble ribosomes during the recovery from growth arrest.

Here, we show that an *E. coli* Δ*rimM* strain recovers from growth arrest markedly more slowly than isogenic Δ*rbfA*, Δ*ksgA* (a late-stage maturation factor for the 30S decoding site)^29^, or Δ*S9* (lacking r-protein uS9) strains (Fig. 1a). The ribosomes of the Δ*rimM* strain are specifically deficient in the translocation step of the elongation cycle, indicating deficiency in the 30S head domain^30–32^. This deficiency provides the basis for over-production of the ribosomal machinery across all phases of the growth cycle. Cryo-EM analysis of ribosomal particles isolated from the early exponential phase of the Δ*rimM* strain revealed the persistent presence of IF1 and IF3 in the 30S subunit, delayed recruitment of r-proteins (uS3, uS10, uS13, uS14, uS19) to the head domain, as well as bound RsfS in both pre-50S and mature 50S particles. The persistent presence of IF3 and RsfS in pre-30S and pre-50S particles, respectively, is also observed in proteomic analysis, supporting the anti-association activity of each^33^ and reinforcing the notion of ribosome quality control that prevents the pre-mature assembly of defective ribosomes. This large-scale ribosome quality control in the absence of RimM, which delays ribosome recovery to allow time to coordinate with multiple factors, highlights the importance of proper 3′-domain folding of the 16S rRNA and proper assembly of the head domain in the bacterial exit from growth arrest.

## Results

### An E. coli ΔrimM strain exhibits a pronounced delay in recovery from growth arrest

To identify the most critical factors for ribosome recovery from growth arrest, we compared an *E. coli* Δ*rimM* (MG1655) strain with strains that lack viable checkpoints of 30S maturation (Fig. 1a). The Δ*rbfA* strain affects the maturation of the 16S rRNA 5′-end and decoding center^14,18^; the Δ*ksgA* strain removes the terminal dimethylation checkpoint at the decoding site^29^; the Δ*rsmE* strain removes the methyl transferase that installs the m^3^U1498 modification in the 16S rRNA^34^ located in helix 44 (h44) near the A-site; the Δ*S9* (Δ*rpsI)* strain alters a core r-protein that shapes the geometry of the head domain of the 30S subunit^35,36^; and the *S13-mut (rpsM-mut)* strain, which lacks the C-terminal residues (82–117) of the uS13 r-protein in the 30S head region, affects the formation of inter-subunit bridges with the 50S subunit^37^. Together these strains provide a spectrum to examine the 30S recovery pathway from early to late stages, enabling side-by-side comparisons.

An *E. coli* wild-type (*wt;* MG1655) strain and a pair of isogenic Δ*rimM* and Δ*rbfA* strains were grown in LB to the stationary phase overnight (> 12 h), diluted into fresh LB, and the growth recovery was monitored by OD_600_ (Fig. 1b). While the *wt* strain had a lag phase of recovery of 1.1 h and Δ*rbfA* had a lag phase of 2.3 h, Δ*rimM* had a much longer lag phase of 4.5 h (Fig. 1b-d). The long lag phase of Δ*rimM* was unexpected and striking, in contrast to the lag phase of Δ*S9* (Δ*rpsI*; 1.1 h), *S13-mut* (*rpsM-mut;* 1.4 h), Δ*ksgA* (1.6 h) and Δ*rsmE* (1.1 h) (Fig. 1e-g). The pronounced lag phase of Δ*rimM* relative to the *wt* was also observed after 5 overnights of growth arrest in LB (Supplementary Fig. 1a-c), and after one overnight of growth arrest in the minimal media M9 (Supplementary Fig. 1d-f) and in the defined media MOPS supplemented with glucose or galactose (Supplementary Fig. 1g-i). Additionally, as cells exited the lag phase and began the growth phase (OD_600_ of 0.2-0.3), the growth of Δ*rimM* remained slow relative to the *wt* and Δ*S9* (Fig. 1d and g). Notably, the slow recovery of Δ*rimM* was not due to a loss of viability relative to the *wt* (Supplementary Fig. 1j), or due to the appearance of heterogeneity of cell populations (Supplementary Fig. 1k). However, Δ*rimM* exhibited a severe cold sensitivity (Supplementary Fig. 1l), consistent with the phenotypes that arise from a deficiency in ribosome assembly^38,39^.

Importantly, the slow recovery of Δ*rimM* from growth arrest was not rescued by the over-expression of *rbfA*, but by the over-expression of *rimM* itself; even over-expression of a variant of *rimM* (*rimM-YY*) was unable to fully rescue the slow recovery (Fig. 1h-j). The *rimM-YY* variant expressed a protein with reduced binding affinity to the 30S subunit, due to alanine substitutions of two highly conserved tyrosine residues (Y106A-Y107A) that affect RimM binding to the r-protein uS19 in the 30S head domain^19^. These results indicate that the slow recovery of Δ*rimM* is exclusively and specifically a consequence of *rimM* deficiency, rather than off-target effects, and that the high affinity of RimM binding to the 30S subunit is a key regulator of ribosome recovery from growth arrest.

To determine how RimM regulates ribosome recovery from growth arrest, we compared the cellular ribosome content between the *wt* and Δ*rimM* strains across three growth phases. Cells were grown to the stationary phase overnight (> 12 h), diluted into fresh LB, and sampled at OD_600_ of 0.1, 0.4, and 1.0, representing the recovery, early exponential, and stationary phases, respectively. The ribosome content was measured using the ratio of total RNA over total protein ratio, which correlates with quantitative mass spectrometry measurements^40^. The results showed that Δ*rimM* produced higher ribosome content than the *wt* across all three phases (Fig. 1k). This suggests an over-compensation effort of *E. coli* to produce more ribosomal subunits and ribosomes to address the deficiency of RimM.

### ΔrimM ribosomes are defective in protein synthesis during the recovery phase

To identify the deficiency of Δ*rimM* ribosomes relative to *wt* ribosomes, we used kinetic assays to assess individual steps of the elongation cycle of protein synthesis. Rather than measuring the overall cellular rate of protein synthesis of a reporter gene^40^, we chose to gather kinetic information to identify the specific steps in an elongation cycle that manifest this ribosomal deficiency. We used our previously established reconstituted system, in which purified 70S ribosomes were programmed with an mRNA and supplied with tRNAs, initiation factors (IF1, IF2, IF3), elongation factors (EF-Tu, EF-G), and GTP^32,41,42^. To correlate with the data above showing ribosomal deficiency of Δ*rimM* across all three phases of a growth cycle, we purified 70S ribosomes from Δ*rimM* and *wt* cells harvested at OD_600_ of 0.1, 0.4, and 1.0, respectively.

We first assayed the dipeptide formation reaction, which evaluated the activity and quality of the ribosome in the A site. An initiation complex containing fMet-tRNA^fMet^ in the P site and an empty A site with a CGU codon was supplied with a ternary complex of EF-Tu, GTP and Arg-tRNA^Arg^. After GTP hydrolysis and Arg-tRNA^Arg^ accommodation, the ribosome synthesized dipeptide fMet-Arg in the A site (Fig. 2a). Using ^35^S-Met in the initiation complex, we measured dipeptide formation by electrophoretic TLC^32,41,42^. The relative plateau reflected the fraction of active Δ*rimM* ribosomes relative to *wt* ribosomes, while the relative *k*_obs_ reported the rate constant of each relative to each other. The results showed that Δ*rimM* ribosomes were deficient relative to *wt* ribosomes across all three phases of growth, exhibiting active fractions at 70-80% and rate constants at 60-70% (Fig. 2b, c and Supplementary Fig. 2a), indicating a defect in the assembly of the A site throughout the growth cycle.

**Figure 2.**
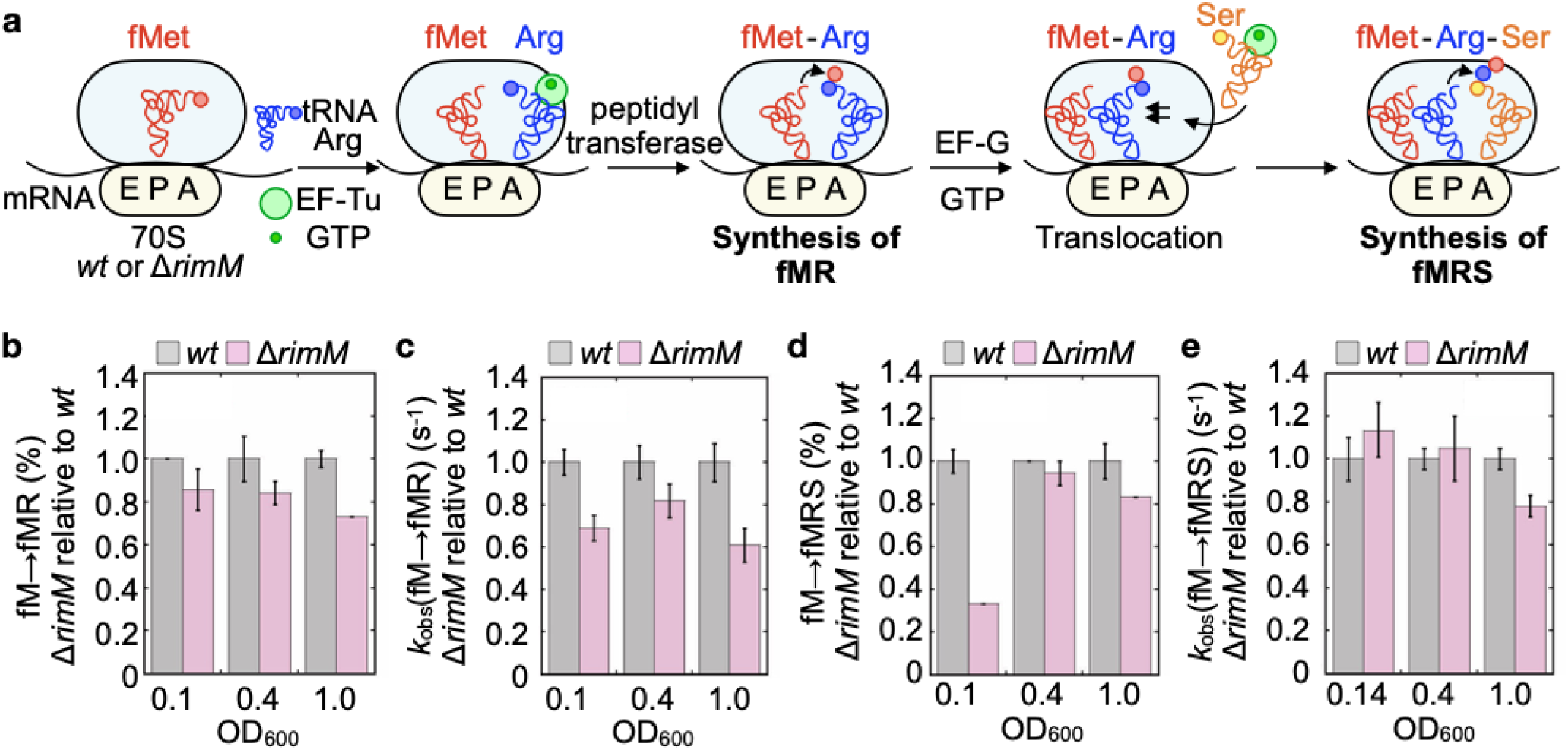
Dipeptide and tripeptide synthesis of *E. coli wt* and Δ*rimM* ribosomes. **(a)** Synthesis of dipeptide (fMet-Arg, fMR) and tripeptide (fMet-Arg-Ser, fMRS) by 70S ribosomes purified from *E. coli wt* and Δ*rimM* strains (MG1655). An initiation complex with fMet-tRNA^fMet^ in the P site, programmed with the coding sequence AUG-CGU-UCG for the first three codons, was mixed with a ternary complex containing Arg-tRNA^Arg^, which promoted peptidyl transfer and the synthesis of fMR. This complex was then mixed with EF-G.GTP to promote translocation, followed by the entry of Ser-tRNA^Ser^ for synthesis of fMRS in the A site. **(b)** Fractional conversion of fM to fMR (%) of Δ*rimM* relative to *wt* ribosomes harvested at indicated OD600’s. **(c)** Rate constant (*k*obs) of synthesis of dipeptide fMR by Δ*rimM* relative to *wt* ribosomes harvested at indicated OD600’s. **(d)** Fractional conversion of fM to fMRS (%) of Δ*rimM* relative to *wt* ribosomes harvested at indicated OD600’s. **(e)** Rate constant (*k*obs) of synthesis of tripeptide fMRS by Δ*rimM* relative to *wt* ribosomes harvested at indicated OD600’s. For panels b-e, error bars, mean ± SD (n = 3).

We next assayed tripeptide formation in the A site, where a dipeptide was synthesized first, followed by translocation of the deacylated tRNA^fMet^ and the dipeptidyl ^35^S-Met-Arg-tRNA^Arg^, after which Ser-tRNA^Ser^ entered the A site, leading to synthesis of the tripeptide ^35^S-fMet-Arg-Ser (Fig. 2a). Kinetic parameters of this tripeptide formation were composite terms, including dipeptide formation in the A site, translocation of the dipeptidyl-tRNA from the A to the P site, and tripeptide formation in the A site. Since the *k*_obs_ of tripeptide formation of both the Δ*rimM* and *wt* ribosomes was consistently lower than the respective dipeptide formation (Supplementary Fig. 2), we concluded that the composite terms of tripeptide formation were rate-limited by the event of translocation. Notably, while the *k*_obs_ of Δ*rimM* ribosomes was similar relative to the respective *wt* ribosomes across all three phases (Fig. 2e), the active fraction of Δ*rimM* ribosomes was particularly low during the recovery phase (Fig. 2d), indicating a specific deficiency of Δ*rimM* ribosomes in the translocation event. As the quality and efficiency of translocation is critically dependent on the 30S head domain^30–32^, the deficiency of Δ*rimM* ribosomes in the recovery phase is consistent with an immaturely assembled head domain. This provides a kinetic basis for ribosome deficiency, particularly the deficiency of the head domain, for the slow recovery of *E. coli* from growth arrest in the absence of RimM.

### IF1 and IF3 safeguard the 30S head domain maturation in ΔrimM strain

To uncover the structural basis for this functional defect (Fig. 2d), we examined ribosomal particles from the Δ*rimM* strain by single-particle cryo-EM, using *E. coli* Δ*rimM* cells collected at OD_600_ = 0.4 (early exponential phase). Here, ribosomes were purified from the *E. coli* MRE600 strain using high-salt sucrose gradient fractionation to preserve maturation intermediates and to isolate only stably ribosome bound auxiliary factors. Notably, the MRE600 strain exhibited the same prolonged recovery of Δ*rimM* cells as in MG1655, with similar ribosome profiles between strains and conditions, including an over-representation of ribosomal subunits relative to 70S ribosomes, which indicated an ongoing process of 30S maturation and 70S assembly (Supplementary Fig. 3).

Cryo-EM analysis of the 30S_Δ_*_rimM_* revealed a fraction of immature 30S particles lacking head domain density (Supplementary Figs. 4 and 5a), consistent with previously described intermediates in 30S biogenesis^4,5^. Notably, we identified three distinct maturation states of the 30S_Δ_*_rimM_* subunit (I-III), all without the presence of the mRNA but bound with initiation factors IF1 and IF3 at their canonical positions near the 30S platform and A site^43,44^, respectively (Fig. 3 and Supplementary Fig. 6). These three states represent progressive stages of head domain formation, according to the Nomura assembly map^45,46^ for the stepwise incorporation of r-proteins into the 30S head domain and the concurrent stabilization of the 3′-major domain of the 16S rRNA (Fig. 3).

**Figure 3.**
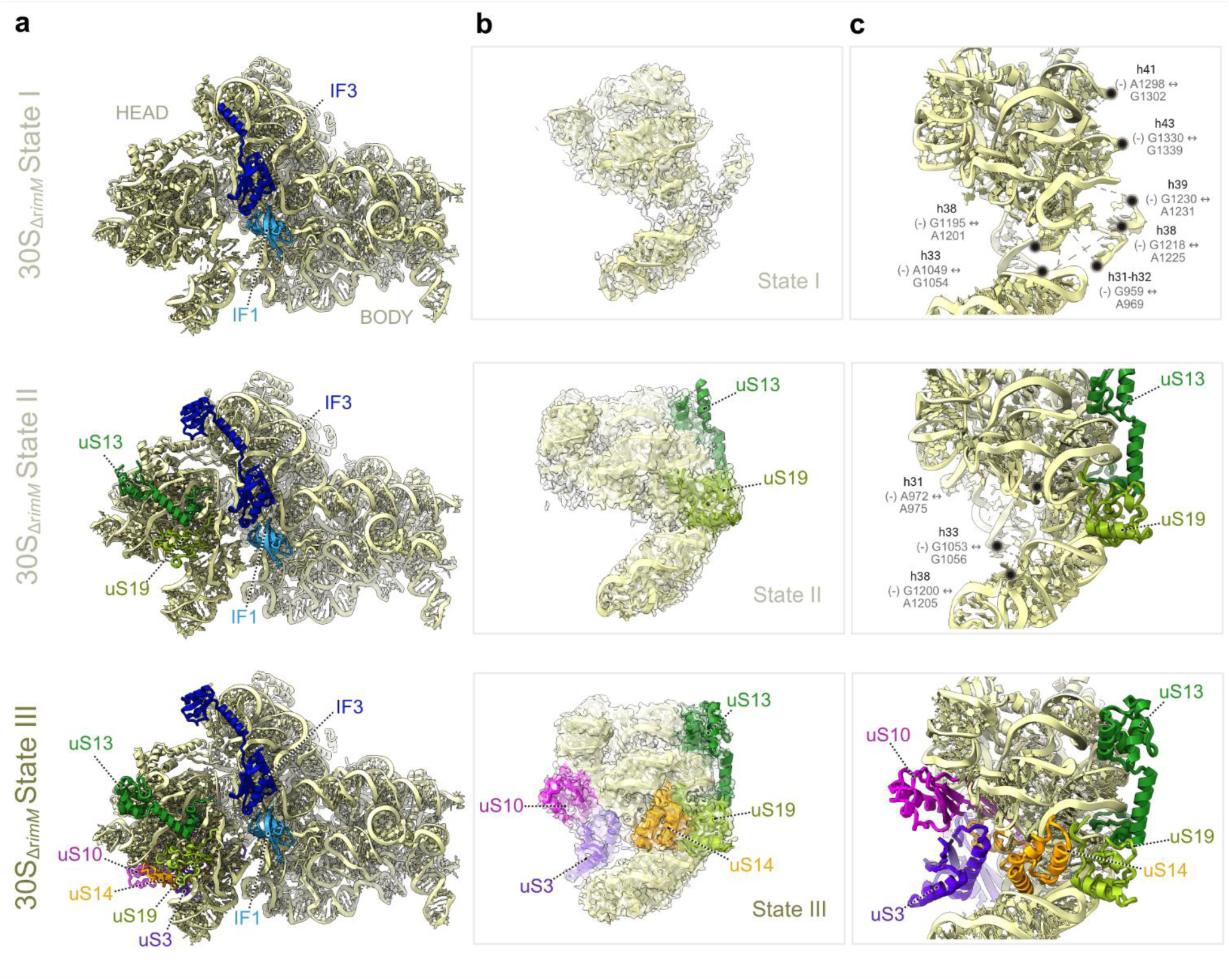
Stepwise maturation of 30S head domain with bound IF1 and IF3 in *ΔrimM* strain. **(a)** Structural models of the 30S subunit (light yellow) representing distinct stages of head domain maturation (top to bottom: state I, state II, and state III), with bound IF1 (light blue) and IF3 (dark blue) in all three states. In state I, the 30S head domain is highly disordered and lacks r-proteins uS3, uS10, uS13, uS14, and uS19. In state II, the incorporation of uS13 (green) and uS19 (lemon) indicates partial head stabilization. In state III, all late-assembling head domain r-proteins - uS3 (purple), uS10 (magenta), and uS14 (orange) - are present, reflecting a mature 30S head conformation. **(b)** Cryo-EM density surface views (non-sharpened original cryo-EM maps) of the 30S head domain combined with structural models across the three 30S maturation states of the head domain, highlighting the progressive incorporation of r-proteins and compaction of the 16S rRNA. The cryo-EM maps are shown at 2-3 σ for states I to III, respectively. **(c)** Close-up views of the 3′-major domain of the 16S rRNA within the head region, showing the gradual stabilization of critical 16S rRNA helices. Disordered or flexible regions of the 16S rRNA, along with their associated nucleotides, are annotated where possible. The coloring of r-proteins and corresponding helices is as in panel (a).

In state I, densities for r-proteins uS3, uS10, uS13, uS14 and uS19 were absent, and the 30S head domain displayed high conformational flexibility. This flexibility was especially pronounced in the 3′-major domain of the 16S rRNA, which normally depends on these r-proteins for stabilization. According to the Nomura assembly map^45,46^, uS13 and uS19 are secondary assembly proteins, while uS3, uS10, and uS14 are part of the tertiary (late) assembly stage. Structural analysis of the 3′-major domain of the 16S rRNA showed disordered regions in several key helices, including nucleotides 952–953 (helix h31), 959–969 (h31–32 junction), 983–989 (h32), 1049–1054 (h33), 1195–1201 and 1218–1225 (h38), 1230–1231 (h39), 1298–1302 (h41), 1307–1309 (h42), and 1330–1339 (h43) (Fig. 3a-c, top row). These helices form the structural core of the 30S head and are essential for intersubunit bridge formation and the accommodation of tRNAs^47,48^. The absence of uS3, uS10, uS13, uS14, and uS19 correlated with destabilization in these regions, reinforcing the critical roles of these r-proteins in structuring the head domain during 30S maturation^5,49^. Therefore, 30S_Δ_*_rimM_* state I represents an early intermediate in the maturation of the head domain in the absence of RimM.

In 30S_Δ_*_rimM_* state II (Fig. 3a-c, middle row), r-proteins uS13 and uS19 (secondary assembly proteins) were bound to the 30S head domain. Their incorporation led to localized stabilization of the 3′-major domain of the 16S rRNA, showing partial compaction and structural ordering of the head domain as it progressed toward full maturation. This stabilization was particularly evident in rRNA helices h41, h42, and h43, which directly interacted with uS13 and uS19. However, several regions of the 16S rRNA within the head domain remained flexible, especially rRNA helices encompassing nucleotides 972–975 (h31), 1053–1056 (h33), and 1200–1205 (h38). Proper folding and stabilization of these regions would depend on the subsequent incorporation of tertiary assembly r-proteins, particularly uS3, uS10, and uS14.

In 30S_Δ_*_rimM_* state III (Fig. 3a-c, bottom row), the late-assembly r-proteins uS3, uS10, and uS14 were fully incorporated into the 30S head domain. The 3′-major domain of the 16S rRNA showed ordered structures in regions that were previously dynamic or unfolded, indicating that structural maturation of the head domain was complete. Structural comparison with the mature 30S subunit^44^ confirmed that state III closely matches the native configuration in both r-protein composition and 16S rRNA folding (Supplementary Fig. 6), whereas states I and II exhibited progressive improvement in these features. IF1 and IF3 remained bound throughout all three states, indicating that their persistent presence plays a regulatory role^33,50^ to stabilize immature 30S subunits and to guide their stepwise maturation. This ensures that only properly assembled 30S subunits proceed to association with the 50S subunit to assemble functional 70S ribosomes for translation.

### RsfS controls the 50S subunit maturation and 70S assembly

To test whether the deficiency of 30S assembly in the Δ*rimM* strain was coordinated with the maturation of the 50S subunit, we performed single-particle cryo-EM on 50S subunits isolated from the *E. coli* Δ*rimM* strain at an OD_600_ of 0.4. The analysis revealed two major populations of 50S particles (Supplementary Fig. 5b and 7). One population consisted of immature 50S (pre-50S) subunits lacking helix 68 (-H68), a late-assembling helix of the 23S rRNA that is inserted between the L1 stalk and the central protuberance during the final stages of 50S maturation^21,24,25^. The second population consisted of fully mature 50S subunits with helix H68 correctly folded and integrated in its canonical position. Notably, some fractions of the pre-50S_Δ_*_rimM_* and 50S_Δ_*_rimM_* particles were bound with the ribosome silencing factor RsfS (Fig. 4). Cryo-EM maps clearly resolved RsfS at its canonical binding site on the intersubunit face of the 50S subunit, similar to previous structural studies of the 50S subunit isolated from *S. aureus*^26^ (Supplementary Fig. 8) and *M. tuberculosis*^51^. RsfS adopted a compact α/β fold and was in direct contact with r-protein uL14 near the L11 stalk. Its binding site formed a stable cleft between uL14 and adjacent 23S rRNA helices, which was supported by a dense network of electrostatic and hydrophobic interactions (Fig. 4). RsfS remained stably bound at this site regardless of the presence or absence of helix H68, which is located distantly on the opposite side of the 50S subunit near the L1 stalk (Fig. 4). These findings indicate that RsfS plays a dual role, acting as a maturation checkpoint that senses the biogenesis state of the 50S subunit^22^, and as an anti-association factor that prevents premature 70S assembly^26^.

**Figure 4.**
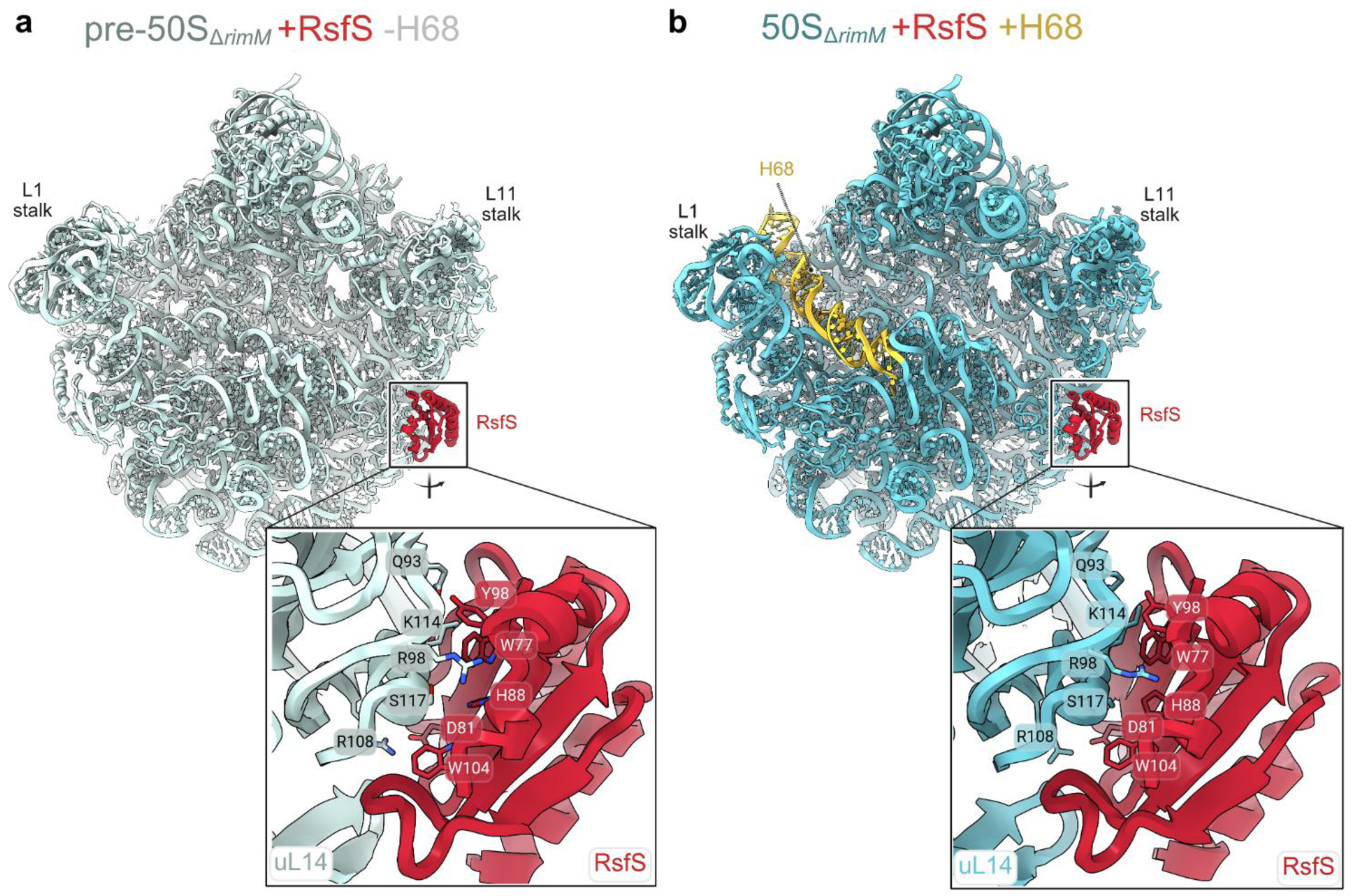
RsfS bound to both pre-50S and 50S ribosomal subunits in Δ*rimM* strain. **(a)** Cryo-EM structure of immature pre-50S–RsfS complex lacking helix H68 (-H68). The 50S ribosomal subunit is shown in light blue in a crown view from the intersubunit interface, with RsfS highlighted in red. The close-up view of the RsfS–uL14 interface shows key interacting residues. **(b)** Cryo-EM structure of mature 50S–RsfS complex showing the presence of the ordered helix H68 (highlighted in yellow). The 50S subunit is shown in cyan in the same view as in (a). The close-up view of the RsfS–uL14 interface shows the interacting residues as in (a).

### IF1, IF3 and RsfS control ribosomal maturation and association of subunits

The occupancy of IF1, IF3 and RsfS in Δ*rimM* ribosomal samples suggested an important role of each in the ribosomal maturation process. To directly assess the importance of each, we performed a quantitative comparison of ribosomal samples from Δ*rimM* relative to the *wt* strain (MRE600) collected at OD_600_ of 0.4 (Supplementary Figs. 4, 5, 7, 9-13). This comparison showed that, in mature particles, the ribosomes of the two strains were structurally indistinguishable, whereas in immature particles, the two strains differed substantially in the occupancy of the factors, particularly in the 30S and 50S subunits (Fig. 5).

**Figure 5.**
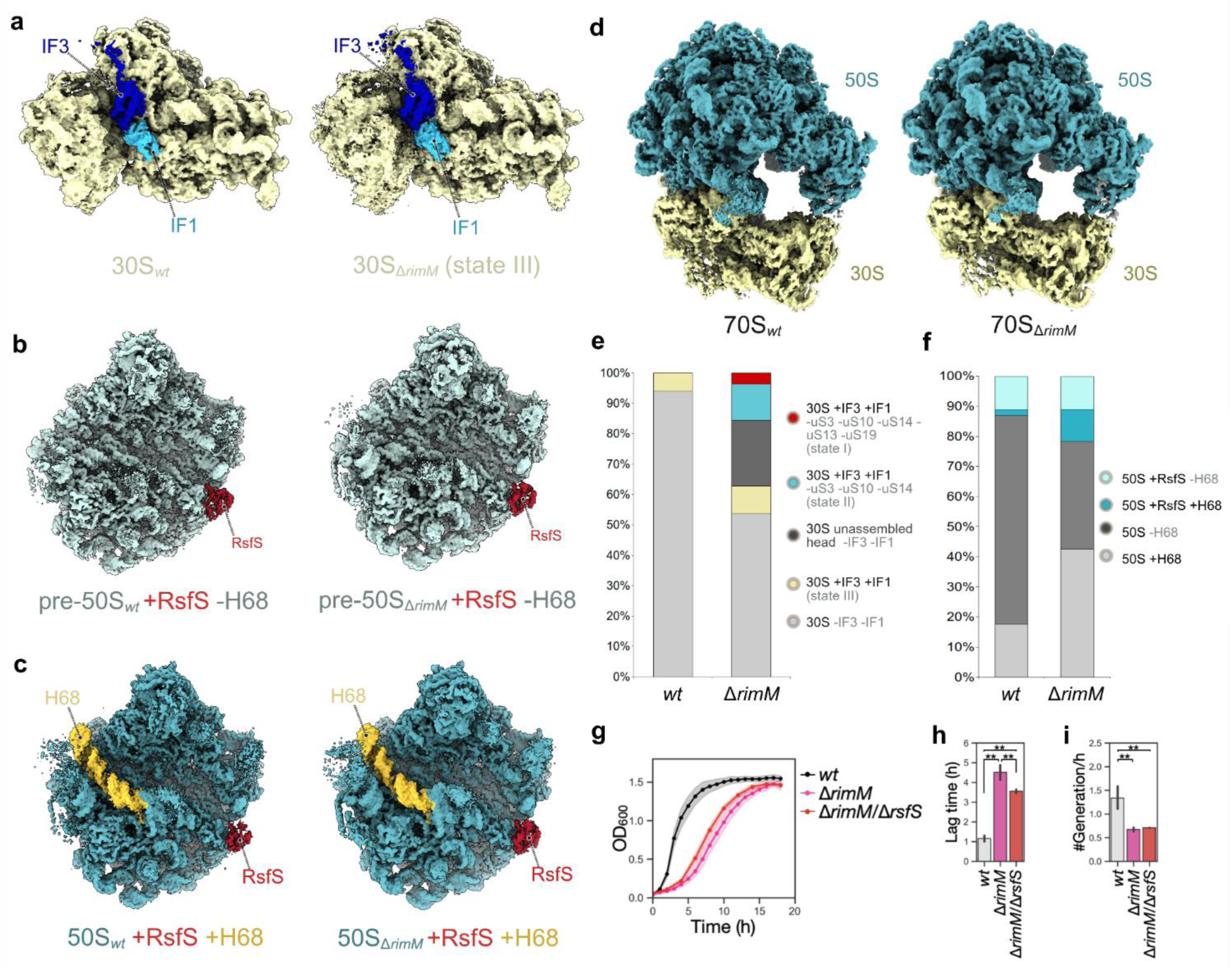
IF1, IF3 and RsfS occupancy in *wt* and Δ*rimM* strains. **(a)** Non-sharpened original cryo-EM maps (shown at 2.5 σ) of mature 30S subunits from the *wt* and Δ*rimM* strains (state III), respectively, showing IF1 (light blue) and IF3 (dark blue) bound to the 30S subunit (light yellow). **(b)** Non-sharpened original cryo-EM maps (shown at 3.5 σ) of pre-50S subunits lacking H68 (–H68) from the *wt* and Δ*rimM* strains showing RsfS (red) bound at the intersubunit interface. **(c)** Non-sharpened original cryo-EM maps (shown at 3.5 σ) of mature 50S subunits in the *wt* and Δ*rimM* strains. RsfS (red) remains bound with H68 (gold) present. **(d)** Non-sharpened original cryo-EM maps (shown at 3.0 σ) of 70S ribosomes from the *wt* and Δ*rimM* strains, composed of mature 30S (yellow) and 50S (cyan) subunits. **(e)** Occupancy analysis of 30S particles in the *wt* and Δ*rimM* strains shown in percentages normalized against the total amounts of 30S particles observed. **(f)** Occupancy analysis of 50S particles in the *wt* and Δ*rimM* strains shown in percentages normalized against the total amounts of 50S particles observed. **(g)** Growth profiles of *wt* (n = 13), Δ*rimM* (n = 13), and Δ*rimM*/Δ*rsfS* (n =3) strains of MG1655 in LB. **(h)** Lag time of each culture in (g). **(i)** Growth rate of each culture in (g). Error bars, mean ± 95% confidence interval. \**p* < 0.05, \*\**p* < 0.01; two-tailed Welch’s t-test.

In the *wt* strain (Fig. 5a), fully assembled head domains of 30S subunits were found without incomplete intermediates. The only IF1/IF3-bound 30S class corresponded to a fully mature subunit, matching 30S_Δ_*_rimM_* state III, in which the 16S rRNA folding, r-protein placement, and IF1/IF3 binding sites were fully in matured state in both the *wt* and Δ*rimM* strains. Key structural domains – including the platform, body, and head – were intact, and the decoding center, comprising helices h44, h45, were fully ordered in both strains. This suggests that, once the head domain maturation is complete, 30S_Δ_*_rimM_* subunits can support translation as efficiently as *wt* subunits.

In contrast, in the Δ*rimM* strain, immature 30S particles represented by 30S_Δ_*_rimM_* states I and II, were accumulated as shown by immature head domains lacking late-binding r-proteins while retaining IF1 and IF3 (Fig. 5e). Notably, the disordered head domain structures agree with those of earlier cryo-EM studies collected from an *rimM*-depleted strain^4,5^. Quantitative cryo-EM revealed (Fig. 5e) that IF1 and IF3 are more abundant in the immature 30S particles (30S_Δ_*_rimM_* I and II) compared to the *wt*. Complementary mass spectrometry showed elevated IF3 levels, while IF1 remained unchanged in the Δ*rimM* strain during the early exponential phase, suggesting that IF3 upregulation is a key regulatory mechanism in response to the absence of RimM (Supplementary Fig. 14).

The comparison of 50S subunits between the *wt* and Δ*rimM* strains revealed no significant differences in the 23S rRNA core or r-protein composition (Fig. 5b, c). Quantitative analysis of 50S particle populations (Fig. 5f) showed that in the *wt* strain, most 50S subunits were either fully mature or in a late-stage assembly intermediate lacking helix 68 (-H68) of the 23S rRNA. Only a small fraction of these particles was bound by RsfS. In contrast, the Δ*rimM* strain exhibited a higher proportion of mature 50S subunits, possibly a response to defective 30S assembly by overproducing the 50S subunits (Fig. 1k and Supplementary Fig. 3c). Notably, the number of RsfS-bound 50S particles was also elevated in Δ*rimM*, including both pre-50S and mature 50S subunits (Fig. 5f). Protein quantification showed increased RsfS levels in the Δ*rimM* strain during the early exponential phase, indicating that RsfS upregulation is a key response to the absence of RimM (Supplementary Fig. 14). Notably, the *ΔrsfS* deletion did not alleviate the prolonged recovery phase observed in the Δ*rimM* strain (Fig. 5g-i), likely suggesting that the high occupancy of IF1 and IF3 that remained bound to the 30S particles (Fig. 5e) was sufficient to prevent subunit joining.

Moreover, we found that 50S subunits of both the Δ*rimM* and *wt* strains contained a second copy of the r-protein bS20. While bS20 is typically located on the body of the 30S subunit, this second copy was observed in the 50S subunit, positioned near helices H4, H18, and H28, where it formed close contacts with the r-protein uL4 (Supplementary Fig. 15a,b). Notably, a recent cryo-EM analysis of *P. urativorans* ribosomes detected bS20 on both small and large subunits^52^, where the position in the large subunit matched that of our 50S maps. This second copy appeared to be a neutral and heterogeneous feature with no clear functional role^52^.

Despite the pronounced differences between the Δ*rimM* and *wt* strains either in the composition or maturation state of 30S and 50S subunits, the 70S ribosomes were identical (Fig. 5d). Structural comparisons revealed no detectable differences between the assembled ribosomes. In both cases, mature 30S and 50S subunits were incorporated without any rRNA folding defects, misalignment of intersubunit bridges, or changes in the positioning of r-proteins. The only notable addition in the 70S ribosomes was the presence of a second copy of the r-protein bS20 in both strains (Supplementary Fig. 15c, d), similar to that observed in isolated 50S subunits. Both the 30S decoding center and the 50S peptidyl transferase center were correctly formed. Thus, while 30S subunits of Δ*rimM* are delayed in maturation, once they are matured, they are readily assembled with mature 50S subunits into 70S ribosomes that have the normal structural architecture.

## Discussion

### RimM is a key determinant of ribosome recovery from growth arrest

Rapid ribosome recovery from growth arrest is important for the proliferation of every cellular organism. While the rRNA and protein components are sufficient for ribosome self-assembly, additional assembly factors are needed to avoid delay. Of the 10-20 assembly factors for maturation of the 30S subunit^36^, RbfA and RimM are both critical for ribosome recovery in *in vivo* experiments, guiding the step-wise organization of the 16S rRNA and recruitment of r-proteins^18^. Between the two, RbfA has long been thought of as the major determinant rather than RimM, not only because of the longer residence time of RbfA on pre-30S particles, but also because of additional considerations. (i) RbfA is a high-abundance protein in *E. coli*, whereas RimM is a low-abundance protein^53^, implicating RbfA in important roles. (ii) While both *E. coli* Δ*rbfA* and Δ*rimM* are cold-sensitive, the Δ*rbfA* strain accumulated substantially more unprocessed 17S rRNA (25%) relative to Δ*rimM* (8%) at the respective non-permissive temperature^54^, indicating a stronger role of RbfA in the maturation of the 30S subunit. (iii) While both Δ*rbfA* and Δ*rimM* accumulated pre-30S particles, 50% of those in Δ*rbfA* were inactive, whereas all fractions in Δ*rimM* were active^54^, indicating a stronger negative effect upon the loss of *rbfA*.

However, despite the above considerations, we show here that the Δ*rimM* strain exhibits a significantly prolonged delay in ribosome recovery relative to the Δ*rbfA* strain in multiple comparisons with other deletion strains (Fig. 1). Additionally, the delay in growth recovery of the Δ*rimM* strain is also significantly longer than *E. coli* strains lacking other checkpoint proteins of 30S maturation (Fig. 1). Further, the delay of Δ*rimM* can only be rescued by RimM itself; not by RbfA, and not by a variant of RimM that has a reduced binding affinity to 30S (Fig. 1h-j). These results establish a specific role of RimM as a key determinant of ribosome recovery from growth arrest, even though the protein binds to pre-30S particles more transiently than RbfA. We suggest that since 30S maturation is driven by progressive structural rearrangements of the 16S rRNA, and since RimM is specifically responsible for the folding of the 3’-major domain of the 16S rRNA localized in the head domain, our findings emphasize that the successful folding of this portion of the 16S rRNA (Fig. 3) is one of the most critical factors for the rapid recovery of a ribosome from growth arrest. The importance of RimM for folding of the 3’-domain of the 16S rRNA is consistent with our observation that Δ*rimM* ribosomes are specifically deficient in the translocation step of protein synthesis (Fig. 2d,e), where the head domain is critically important^30–32^, and that this deficiency is most pronounced during the recovery phase, where the assembly of the head domain is delayed.

Notably, assembly of the 3’-domain of the 16S rRNA is a challenging task in ribosome biogenesis, requiring a substantial energy investment. *In vitro* studies showed that the rRNA must undergo a heat-dependent rearrangement for r-proteins uS2, uS3, uS10, uS14, and uS21 to bind^55^, of which uS3, uS10, and uS14 are missing in the Δ*rimM* pre-30S particles. One possibility is that the high-energy barrier of the 3’-domain assembly is created by an intrinsic rRNA folding intermediate that is resolved by RimM-mediated rearrangements, and that the loss of RimM causes the accumulation of immature assembly complexes that prevent the recruitment of specific r-proteins. Alternatively, the high-energy barrier may arise from a subset of r-proteins that bind to the 16S rRNA before uS3, uS10, and uS14. These proteins could induce rRNA rearrangements that form a structural trap requiring RimM for resolution. Thus, RimM is an important factor in the energy landscape of the 3’-domain assembly of the 16S rRNA. Although the energy landscape of the Δ*rimM* strain cannot be identical to that of the *wt* strain, which expresses RimM, this study emphasizes the importance of the 3’-domain assembly of the 16S rRNA in bacterial recovery from growth arrest.

### IF1, IF3 and RsfS coordinate to regulate ribosome recovery in the absence of RimM

We show here that *E. coli* cells, upon the loss of RimM, launch a large-scale coordinated program to delay ribosome biogenesis to provide time for self-assembly to ensure the fidelity of protein synthesis. This coordinated program involves the accumulation of IF1 and IF3 on the 30S subunit, and the accumulation of RsfS on the 50S subunit, to enable the delayed incorporation of uS3, uS10, uS13, uS14, and uS19 into the head domain. This delay is the underlying basis of the substantially delayed ribosome recovery time after growth arrest (Supplementary Fig. 3a). It is also the basis for the observation that, despite a delay that allows self-assembly of the 3′-domain of the 16S rRNA, Δ*rimM* ribosomes eventually become fully functional to support growth to the same densities as *wt* ribosomes under various conditions (Fig. 1b–h and Supplementary Fig. 1).

We summarize our results in a model to illustrate the coordination of IF1, IF3, and RsfS to prevent premature ribosome assembly in Δ*rimM* (Figure 6). In *wt* cells, RimM facilitates proper 3’-domain folding of the 16S rRNA and stabilizes rRNA–protein interactions, enabling the timely and transient binding of IF1 and IF3 to prepare the 30S subunit for translation initiation. This enables the efficient binding of IF2-fMet-tRNA^fMet^ and subunit joining to transition to the formation of an active 70S ribosome^44,56^. In Δ*rimM* cells, in contrast, the structural defects are primarily associated with the 30S subunit (Fig. 3), but do not compromise the formation of the 50S or the 70S ribosome (Fig. 5b-d). As such, the defective 30S particles retain IF1 and IF3 longer to allow time for self-assembly of the 30S subunit, while the 50S particles recruit higher levels of RsfS to prevent premature subunit joining. The coordination of the three factors ensures that only properly matured subunits are engaged in translation. Notably, while the assembled 70S ribosomes of Δ*rimM* cells are identical in structures to those of *wt* cells, we observed kinetic deficiencies of these ribosomes in protein synthesis (Fig. 2), indicating that a fraction of the self-assembled ribosomes in Δ*rimM* cells could be kinetically or dynamically deficient. It is possible that the self-assembly process, in the absence of RimM, would take an alternative route than the factor-assisted route.

**Figure 6.**
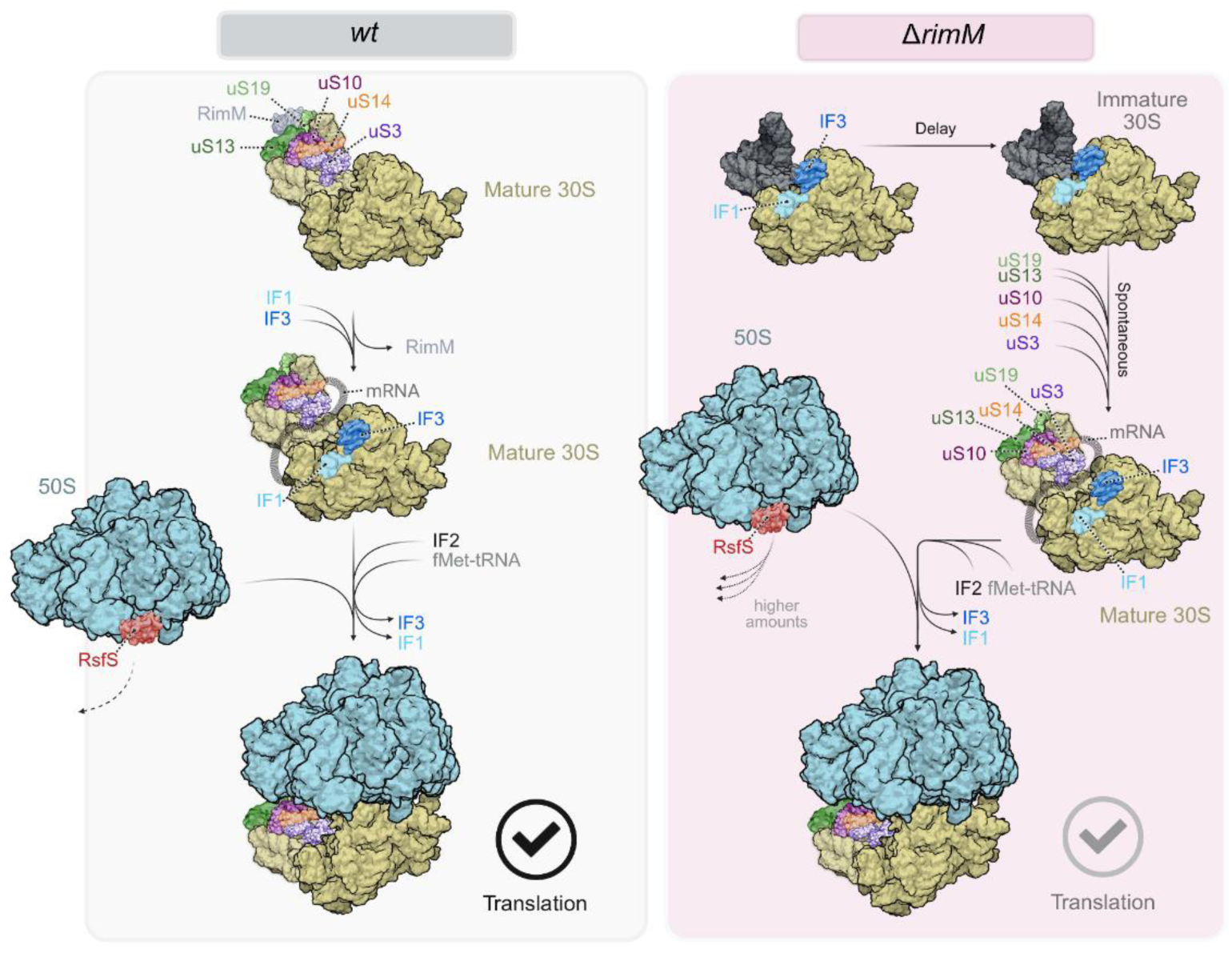
A model for 70S assembly and translation recovery. Left: In *wt* cells, RimM enhances recruitment of 30S head ribosomal r-proteins (uS3, uS10, uS13, uS14, uS19) and facilitates correct 30S subunit maturation. IF1 and IF3 interact with mature subunits, allowing efficient translation initiation after following subunit joining. Right: in Δ*rimM* cells, IF and IF3 delay 30S head maturation and prevent subunit joining. This delay allows time for slow-joining of r-proteins to be progressively incorporated into the head domain. Upregulated RsfS levels safeguard the 50S subunits until 30S assembly is complete. Once the assembly of the 30S head domain is complete, the subunit joining takes place to resume protein synthesis.

The involvement of IF1 and IF3 in ribosome biogenesis in Δ*rimM* suggests a second role of these factors beyond their canonical involvement in the initiation of protein synthesis^43^. In the second role, they would help screen against immature subunits from entering protein synthesis. This quality control by IF1 and IF3 would be in line with emerging models in which IF3 acts as a checkpoint factor, preventing premature 50S joining or aberrant translation initiation by structurally defective 30S subunits^14,50^. Additionally, while RsfS typically binds to pre-50S subunits as an assembly factor^22^, its presence on the pre-50S and mature 50S subunits in our study indicates a specific role in the absence of RimM. We suggest that, since the binding site of RsfS in pre-50S particles does not overlap with those of translation factors or antibiotics, the purpose of this binding is to allow the stabilization of helix H68 of the 23S rRNA, which is a critical component of the 50S subunit that determines the quality of the translocation step^57^. However, in the absence of RimM, we suggest that RsfS has a unique role of “sensing” the accumulation of pre-30S particles and it is pre-emptively binding to the 50S subunit to forestall premature subunit joining^26^. Importantly, our *E. coli* findings possibly point to a broader and conserved mechanism of safeguarding against the premature joining of ribosomal subunits, using initiation and anti-association factors as the checkpoints. In mitochondria, the MALSU1 factor (the counterpart of bacterial RsfS) similarly blocks subunit joining during the late stage of 60S subunit assembly^58,59^. This, together with the report that mtIF3 remains bound to pre-40S particles during late assembly^60,61^ and our observations reported here, suggests a similar role of RsfS and IF3 to screen against defective subunits from joining. This model may represent a common design principle that reflects an ancient solution for preserving ribosome fidelity during recovery from stress.

Finally, we observed an increase in ribosomal content in Δ*rimM* cells across all phases of growth (Fig. 1k), indicating that *E. coli* cells change their cellular resources to generate more of the ribosome apparatus. The energetic costs associated with the synthesis and assembly of the ribosome apparatus far exceed those for the up-regulation of a few specific proteins (e.g. IF3 and RsfS; Supplementary Fig. 14), indicating a large-scale programmatic change in energy allocation to prevent further delays in growth recovery. The prolonged ribosome recovery in Δ*rimM* cells relative to *E. coli* cells lacking other ribosome biogenesis factors (Fig. 1b,e) suggests a uniquely low adaptability of Δ*rimM* in response to environmental changes, based on the inverse relationship of adaptability and the lag time^62^. Indeed, Δ*rimM* cells exhibit a strong sensitivity to cold temperature (Supplementary Fig. 1l). The low adaptability, and the low energy state of the cells, raise the possibility of targeting RimM in a new anti-bacterial strategy that should severely compromise bacterial viability in the presence of additional environmental stress, such as antibiotics. This anti-RimM strategy warrants favorable consideration, as RimM is conserved across a wide range of bacterial species, from fast-growing proteobacteria to slow-growing actinobacteria. In contrast, with the exception of chloroplast^63^, eukaryotes have no RimM homologues and have evolved a different strategy for ribosome biogenesis, suggesting the possibility to specifically target bacterial RimM-dependent stress recovery without affecting eukaryotes.

## Methods

### Strain constructions

Isogenic *E. coli* MG1655 strains of Δ*rimM*, Δ*rbfA* and Δ*ksgA* carrying a kanamycin resistance marker (*kan*^R^) were obtained from the Keio collection^64^ and infected into MG1655 recipients. MG1655 strains Δ*rsfS* and Δ*rsmE* were constructed using the λ-Red recombinase system^65^. A kanamycin resistance marker flanked by sequences homologous to the 5′ and 3′ ends of the chromosomal *rsfS* gene and *rsmE* gene were PCR amplified from a pKD4 plasmid using the primers described in Supplementary Table 1. The PCR amplified sequences were electroporated into *E. coli* MG1655 cells carrying the λ-Red recombinase expression plasmid pKD46. Kanamycin-resistant colonies were screened for their correct insertion to the *rsfS* locus and to the *rsmE* locus and were confirmed by PCR using the primers described in Supplementary Table 1. Each gene knockout locus was transferred to the *E. coli* MG1655 strain by P1 transduction using P1 phage lysates prepared from the corresponding knockout strains. The P1 phage lysate prepared from MG1655 Δ*rimM* was used to generate the MG1655 Δ*rimM*/Δ*rsfS* and MRE600 Δ*rimM* strains. All MG1655 strains were λDE3-lysogenized to enable protein expression from T7 expression vectors with a λDE3 Lysogenization Kit (#69734-M, Novagen). The K-12 strains Δ*rpsI* (lacking the uS9 protein) and *rpsM-*mut (encoding a mutant of uS13 lacking residues 82-117) were obtained from Kurt Fredrick^66^.

### Plasmid constructions

A plasmid expressing His-*rimM,* codon-optimized for *E. coli,* was kindly provided by Dr. Ortega at McGill University. To generate a plasmid expressing His-*rimM*-Y106A/Y107A, site-directed mutagenesis was performed by inverse PCR with primers containing the mutation described in Supplementary Table 2, using pET15b-His-*rimM* as the template. To construct plasmids expressing *rbfA*, the *rbfA* gene was amplified from *E. coli* MG1655 genomic DNA using the primers described in Supplementary Table 2 and cloned into the pET15b vector by restriction enzyme digestion with NcoI and BamHI and ligation with T4 DNA ligase (#M0202, NEB). All constructs were confirmed by Sanger sequencing. An empty vector pET22b was used as a control.

### Growth assay

Cells were inoculated into fresh LB medium, MOPS-rich medium (#M2105, Teknova), or M9 minimal medium at a 1:100 dilution after 1-day or 5-day pre-culture in LB and incubated in 24-well plates at 37°C. Strains carrying a kanamycin resistance marker were grown in media supplemented with kanamycin. OD_600_ was measured over time in an Infinite M200 PRO plate reader (Tecan). MOPS medium was supplemented with glucose (#G0520, Teknova) or with galactose at 0.2% (w/v). The measured OD_600_ was fitted to the logarithmic growth equation:

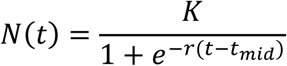

where *N(t)* is the OD_600_ at time *t, K* is the carrying capacity (i.e., maximum OD_600_), *r* is the intrinsic growth rate, and *t*_mid_ is the time when the OD_600_ reaches half of *K*. Although the standard logarithmic equation uses the initial OD_600_ value (*N(0)*) as a parameter ^67^, in this study *t*_mid_ was used, as it had a better fit to the experimental data. Fitting was performed using non-linear least squares with Python v3.7.12 (scipy.optimize.curve_fit). OD_600_ values were baseline-corrected by subtracting the absorbance of the medium-only control. Lag time was defined as the time at which the OD_600_ reaches 0.2, which was calculated using the equation:

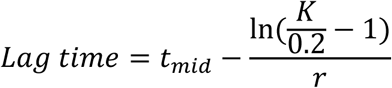

The number of generations per hour was calculated using the equation:

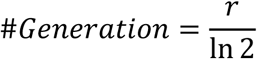

### Ribosome content

Ribosome content was defined as the ratio of [total RNA]/[total protein] for a culture of cells^40^. For total protein quantification, cell cultures of 15 mL, 5 mL, and 2 mL grown at 37°C were pelleted at OD_600_ of 0.1, 0.4, and 1.0, respectively. Cells were washed with water, resuspended in 0.2 mL of water, and flash frozen in liquid nitrogen. The suspension was thawed, lysed with 0.1 mL of 3 M NaOH at 100°C for 5 min, and cooled to room temperature. Protein yield was determined by Bradford assay using Protein Assay Dye Reagent Concentrate (#5000006, Bio-Rad) and measured at 595 nm in an Infinite M200 PRO plate reader (Tecan). For total RNA quantification, the same volumes of culture as for protein quantification were pelleted and flash-frozen in liquid nitrogen. Total RNA was extracted using TRIzol reagent (#15596018, Invitrogen) according to the manufacturer’s instructions and quantified by OD_260_ reading in a Nanodrop spectrophotometer.

### Viability assay

MG1655 strains *wt* and Δ*rimM* were cultured in LB at 37°C for either 1 day or 5 days. To normalize the cell number, both cultures were first diluted to OD_600_ 1.0 with fresh LB, then further diluted 10^5^-fold. A 10 µL aliquot of the diluted culture was spread on LB plates and incubated at 37°C until colonies appeared. Cell counts were calculated from the colony number within six areas of the same size on each plate.

### Cold-sensitivity assay

MG1655 strains *wt* and Δ*rimM* were cultured in LB at 37°C overnight, diluted 1:100 into fresh M9 minimal medium, and grown at 37°C until OD_600_ reached 0.3-0.4 in the exponential phase. Ten-fold serial dilutions (10^0^–10^-5^) were prepared and spotted on M9 plates, which were then incubated at either 37°C or room temperature for 1 to 4 days.

### Ribosome purification for in vitro translation assay

Tight-coupled 70S ribosomes were isolated from *E. coli wt* MRE600_(rif)_ and Δ*rimM* MRE600_(rif+kan)_ cells^68^. Cells were grown in LB medium, harvested at OD_600_ of 0.1, 0.4, and 1.0, flash frozen in liquid nitrogen and stored at −80°C. For ribosome preparation, frozen cells were thawed, washed once in a resuspension buffer (20 mM Tris-HCl, pH 7.5, 30 mM NH_4_Cl, 6 mM Mg(OAc)_2_ and 4 mM βME) and collected by centrifugation at 10,000 rpm at 4°C for 15 min in a SS-34 rotor (Sorvall). The cell pellet was resuspended in fresh resuspension buffer and lysed by three to five passes through a cell disruptor (EmulsiFlex C3, Avestin) at 5,000 psi. Cell debris was removed by two rounds of centrifugation at 16,000 rpm for 30 min at 4°C in a SS-34 rotor. Ribosomes in the clarified supernatant were pelleted through an equal volume of 1.1 M sucrose cushion in buffer A (20 mM Tris-HCl, pH 7.5, 500 mM NH_4_Cl, 10.5 mM Mg(OAc)_2_, 4 mM βME, and 0.5 mM EDTA) by ultracentrifugation at 35,000 rpm for 20 h at 4°C in a 70 Ti rotor (Beckman). The crude ribosomal pellet was resuspended in fresh buffer A and subjected to a second round of pelleting through a sucrose cushion. The pelleted ribosomes were resuspended in buffer B (20 mM Tris-HCl, pH 7.5, 60 mM NH_4_Cl, 10.5 mM Mg(OAc)_2_, 4 mM βME and 0.5 mM EDTA) and separated on a 10-30% sucrose gradient in the same buffer. Gradients were centrifuged at 18,000 rpm for 18 h at 4°C in a SW28 rotor (Beckman). The 70S peak was collected, pooled, and pelleted at 23,000 rpm for more than 24 h at 4°C in a 70Ti rotor. The resulting ribosome pellet was resuspended in buffer B, clarified by centrifugation at 13,500 rpm for 15 min at 4°C in a FA-45-30-11 rotor (Eppendorf), aliquoted, and flash frozen in liquid nitrogen. The ribosome concentration was determined by OD_260_, using 24 pmol per A_260_ unit as the conversion factor.

### In vitro translation assays

Di- and tripeptide formation assays were performed using the codon-walk approach as previously described^32,42^ in a buffer (50 mM Tris-HCl, pH 7.5, 70 mM NH_4_Cl, 30 mM KCl, 3.5 mM MgCl_2_, 1 mM DTT, 0.5 mM spermidine) at 20°C unless otherwise indicated. The three tRNAs used in these experiments were over-expressed in *E. coli* and isolated as part of a total tRNA pool. Each elongator tRNA was separately charged with its cognate aminoacyl-tRNA synthetase. The initiator tRNA^fMet^ was charged with [^35^S]-methionine and formylated with the methyl donor 10-formyltetrahydro-folate by adding formyl transferase to the aminoacylation reaction. All tRNAs were stored in 25 mM NaOAc (pH 5.0) at –70°C. The mRNA template was prepared by *in vitro* transcription and contained a Shine-Dalgarno sequence and an AUG start site: 5’GGGAAGGAGGUAAAA<u>AUGCGUUCG</u>AAG(CCA)_10_, where the first three codons are underlined.

To monitor peptide bond formation, a 70S initiation complex was formed by incubating 70S ribosomes, fMet-tRNA^fMet^, mRNA, and initiation factors IF1, IF2, and IF3 in the HiFi buffer (50 mM Tris-HCl, pH 7.5, 70 mM NH_4_Cl, 30 mM KCl, 3.5 mM MgCl_2_, 1 mM DTT, 0.5 mM spermidine, and 1 mM GTP) for 25 min at 37°C. For dipeptide formation, the ternary complex of Arg-tRNA^Arg^ was mixed with the initiation complex, while for tripeptide formation, both the ternary complex of Arg-tRNA^Arg^ and of Ser-tRNA^Ser^ were mixed with the initiation complex at the same time. Each ternary complex was formed by first incubating EF-Tu in the HiFi buffer for 15 min at 37°C, followed by 15 min in an ice bath after adding a charged tRNA. The complete reaction was formed by rapidly mixing these two solutions in a Kintek model RQF-3 chemical quench apparatus at 20°C. Assays for dipeptide formation contained 0.4 µM 70S ribosome, 0.5 µM mRNA, 0.5 µM [^35^S]-fMet-tRNA^fMet^, 0.5 µM each IF1, IF2, and IF3, 0.5 µM Arg-tRNA^Arg^ (anticodon ACG, where A is modified to I (inosine) in cells), and 0.75 µM EF-Tu in the HiFi buffer. Assays for tripeptide formation additionally contained 0.5 µM EF-G, 0.5 µM Ser-tRNA^Ser^ (anticodon GCU), and 1.5 µM EF-Tu. Timed aliquots from each reaction were quenched with 170 mM KOH and the tRNA-peptidyl linkages were hydrolyzed by incubation at 37°C for at least 15 min. Aliquots of 0.65 µL were spotted onto a cellulose-backed plastic TLC sheet and electrophoresed at 1000 V in PYRAC buffer (62 mM pyridine, 3.48 M acetic acid, pH 2.7) until the marker dye bromophenol blue reached the water-oil interface at the anode^32,42^. The position of the origin was adjusted to maximize separation of the expected oligopeptide products. The separation of unreacted [^35^S]-fMet and each of the [^35^S]-fMet-peptide products was visualized by phosphorimaging and quantified using ImageQuant (GE Healthcare), and kinetic plots were fitted using Kaleidagraph (Synergy software).

### Polysome profiling

MG1655 and MRE600 *wt* and Δ*rimM* strains were cultured in LB at 37°C overnight, diluted 1:100 into fresh LB, and grown at 37°C until OD_600_ reached 0.4 in the early exponential phase. The Δ*rimM* strain carrying a kanamycin resistance marker was grown in LB supplemented with kanamycin. Cell cultures of 50 mL were collected and centrifuged at 7,000 rpm for 10 min at 4°C, and the pellets were frozen in liquid nitrogen. Pellets were thawed and resuspended in 650 µL of lysis buffer (25 mM HEPES, pH 7.5, 100 mM KOAc, 15 mM Mg(OAc)_2_, 1 mM DTT)^69^ supplemented with 6.5 µL of DNase I (#M0303L, NEB). Cells were lysed with 400 mg of 0.1 mm Zirconia/Silica Beads (#11079101z, BioSpec) in at FastPrep-24 5G bead beating (#116005500, MP Biomedicals) instrument with a CoolPrep adapter (#116002528, MP Biomedicals) at 6.5 m/sec for 60 sec, repeated 3 times. Cell lysates were cleared by centrifugation at 13,000 rpm for 10 min at 4°C in a refrigerated microcentrifuge (#C2500-R, Labnet Prism). The RNA concentrations were determined by OD_260_ in a Nanodrop spectrophotometer. Then 25 OD_260_ units of cell lysates were loaded onto a 10-40% sucrose gradient prepared in the lysis buffer. Ribosomes were fractionated by ultracentrifugation at 36,600 rpm for 3.5 hours at 4°C in a SW-41 Ti rotor (Beckman), and the gradient profiles were recorded with a Gradient Station Fractionator (Biocomp).

### LC-MS/MS analysis

Cells were grown under the same conditions as those used for cryo-EM sample preparation. For each biological replicate, 45 mL of culture was harvested by centrifugation at 5,000 rpm for 45 minutes at 4°C (JLA 8.1000 rotor). Protein samples from *E. coli* cell pellets of MRE600 *wt*_(rif)_ and Δ*rimM*_(rif+kan)_ strains (in 3 biological replicates) were extracted in SDT buffer (100 mM Tris-HCl, pH 7.6, 4% SDS, 100 mM DTT) in a thermomixer (Eppendorf ThermoMixer, 15min, 95°C, 750 rpm). After that, all samples were centrifuged (15 min, 20,000 x g) and the supernatants (∼75 μg of total protein) were used for filter-aided sample preparation (FASP) as previously described^70^ using 0.75 μg of trypsin (sequencing grade; Sigma-Aldrich). The resulting peptides were extracted into LC-MS vials with 2.5% formic acid (FA) in 50% acetonitrile (ACN) and 100% ACN with the addition of polyethylene glycol (final concentration 0.001%) and concentrated in a SpeedVac concentrator (Thermo Fisher Scientific).

LC-MS/MS analyses of all peptides were done using the UltiMate 3000 RSLCnano system (Thermo Fisher Scientific) connected to timsTOF HT mass spectrometer (Bruker). Before LC separation, tryptic digests were online concentrated and desalted using a trapping column (Acclaim PepMap Neo C18, dimensions 300 μm ID, 5 mm long, 5 μm particles, 100A, Thermo Fisher Scientific). The trapping column was then washed with 0.1% TFA and the peptides were eluted in backflush mode from the trapping column onto an analytical column (Aurora C18, 75μm ID, 250 mm long, 1.7 μm particles, Ion Opticks) over 90 min in a gradient program (flow rate 200 nL.min^-1^, 3-42% of mobile phase B; mobile phase A: 0.1% FA in water; mobile phase B: 0.1% FA in 80% ACN), followed by a system wash using 80% of mobile phase B. Equilibration of the trapping column and the analytical column was done before sample injection to a sample loop. The analytical column was installed into a Captive Spray ion source (Bruker; temperatures set to 50 °C) according to the manufacturer’s instructions. Spray voltage and dry gas were set to 1.4kV and 3 L.min^-1^, respectively.

MS data were acquired in data-independent acquisition (DIA) mode with the base method m/z range of 100-1700 and 1/k0 range of 0.6-1.4 V×s×cm^-2^. The enclosed diaParameters.txt file defined a m/z 400-1000 precursor range with an equal windows size of 26 Th using two steps for each PASEF scan and a cycle time of 100 ms locked to 100% duty cycle. DIA data were processed in DIA-NN (version 2.1.0)^71^ in library-free mode against the iRT+trypsin database (12 sequences in total, version 2024/10), the common contaminant database cRAP (based on http://www.thegpm.org/crap; 111 sequences in total), and the protein database for *Escherichia coli* (https://ftp.uniprot.org/pub/databases/uniprot/current_release/knowledgebase/reference_proteomes/Bacteria/UP000000625_83333.fasta.gz, version 2025/06, number of protein sequences: 4,402). No optional modification was set during the library preparation, but carbamidomethylation was set as a fixed modification and also trypsin/P enzyme with one allowed missed cleavage and a peptide length of 7-30. The false discovery rate (FDR) control was set to 1% FDR. MS1 and MS2 accuracies as well as scan window parameters were set based on the initial test searches (median value from all samples ascertained parameter values). MBR was switched on.

Protein intensities reported in the DIA-NN main report file were further processed using the software container environment (https://github.com/OmicsWorkflows), version 4.7.7a. The processing workflow is available upon request. Briefly, it consisted of: a) the removal of low-quality precursors and contaminant protein groups, b) the normalization of precursor intensities using the loessF algorithm, c) the replacement of missing values with one or more specific values at the precursor level, d) the calculation of protein group MaxLFQ intensities and transformation of these intensities into log2, and e) differential expression analysis using the LIMMA statistical test. In parallel, data normalization to specific proteins (P0A7V8 and P60438) was also performed. Mass spectrometry proteomics data were deposited to the ProteomeXchange Consortium via PRIDE^72^ partner repository under dataset identifier PXD068732.

### Isolation of 30S, 50S, and 70S ribosomes for cryo-EM

Ribosomal samples were purified from *E. coli* MRE600 *wt*_(rif)_ and Δ*rimM*_(rif+kan)_ strains as previously described^73^. Briefly, cells were grown in LB medium at 37°C and antibiotics were only added to the overnight cultures to maintain selection: rifampicin (0.5 mg/mL) for both *wt* and Δ*rimM* strains, and additionally kanamycin (50 µg/mL) to the Δ*rimM* strain. The next day, the antibiotic-free LB was inoculated with an overnight culture and cells were grown to the early exponential phase (OD_600_ ≈ 0.4). Cells were harvested by centrifugation at 5,000 rpm for 25 minutes at 4 °C (JLA 8.1000 rotor) and washed in ice-cold buffer A (20 mM Tris-HCl, pH 7.5, 100 mM NH_4_Cl, 10 mM MgCl_2_, 0.5 mM EDTA, 6 mM βME). Afterwards, the cells were lysed and homogenized in a Microfluidics M-110P system in buffer B (20 mM Tris-HCl, pH 7.5, 100 mM NH_4_Cl, 12.5 mM MgCl_2_, 0.5 mM EDTA, 6 mM βME). The lysate was clarified by centrifugation at 15,000 rpm for 1 h at 4°C (JA-17 rotor), followed by filtration through a 0.45 μm membrane to remove residual debris. The clarified lysate was layered onto a 37.7% sucrose cushion prepared in buffer C (20 mM Tris-HCl, pH 7.5, 500 mM NH_4_Cl, 15 mM MgCl_2_, 0.5 mM EDTA, 6 mM βME) and ultracentrifuged at 43,000 rpm for 16 h at 4°C (Ti45 rotor). The ribosomal pellet was washed with buffer B on ice and centrifuged at 4,500 rpm for 5 minutes. After the adjustment of the NH_4_Cl concentration to 500 mM in 40 ml buffer C, the sample was centrifuged at 55,000 rpm for 2 hours at 4°C (Ti70 rotor). The resulting pellet was resuspended in 1 ml of buffer B and kept on ice. To isolate the ribosomal samples (30S, 50S and 70S), the resulting ribosomal pellet was layered onto a 10–35% sucrose gradient (in buffer B) and centrifuged at 20,000 rpm for 13 hours at 4°C (SW32 Ti rotor). The fractions of 30S, 50S and 70S were individually collected and concentrated in buffer B (20 mM Tris-HCl, pH 7.5, 100 mM NH_4_Cl, 12.5 mM MgCl_2_, 0.5 mM EDTA, 6 mM βME), flash-frozen in liquid nitrogen, and stored at −80°C.

### Cryo-EM grid preparation, data collection and image processing

Quantifoil R2/1, 300-mesh grids were glow-discharged for 40 s at ∼35 W (30–40 W range) using a Gatan Solarus II. Aliquots of 3.5 μL of 30S, 50S, or 70S samples were applied at a final concentration of 1 μM in buffer A (20 mM Tris-HCl, pH 7.5, 100 mM NH_4_Cl, 10 mM MgCl_2_, 0.5 mM EDTA, 6 mM βME) on the respective grids. Grids were blotted in a Vitrobot Mark IV (Thermo Fisher Scientific) at 4.5°C and ∼95% relative humidity with blot force 0 for 5 seconds, then plunge-frozen in liquid ethane.

All datasets were collected on a Titan Krios (300 kV) equipped with a K3 direct detector and a BioQuantum energy filter (10 eV slit; Gatan). Data were acquired in SerialEM^74^ at 105,000× nominal magnification (pixel size 0.834 Å) with a defocus range of −0.5 to −2.0 µm. Movies were recorded as 40-frame stacks over 2 s (total dose 40 e⁻/Å²) with continuous streaming. MotionCor2^75^ was used for motion correction and frame averaging. Defocus estimation (CTFFIND4)^76^ and particle picking were performed in cisTEM (v1.0.0-beta)^77^. Movies showing ice contamination or excessive drift were excluded based on CTFFIND4 power spectra.

For the 30S datasets, 6,444 micrographs (Δ*rimM*) and 10,995 micrographs (*wt*) yielded 767,668 and 1,523,340 particles, respectively. 2D classification into 50 classes in cisTEM retained 455,076 (Δ*rimM*) and 826,581 (*wt)* 30S particles. For the 50S datasets, 11,367 (Δ*rimM*) and 9,710 (*wt*) micrographs produced 1,523,340 and 1,469,122 particles, with 652,894 (Δ*rimM*) and 970,120 (*wt)* 50S particles retained after 2D classification into 50 classes. For the 70S datasets, 15,893 (Δ*rimM*) and 12,450 (*wt*) micrographs yielded 2,271,232 and 1,579,782 particles, of which 900,281 (Δ*rimM*) and 978,750 (*wt*) 70S particles were retained after 2D classification into 50 classes. Parameter files and particle stacks were then exported from cisTEM, and 4×, 2×, and 1× binned stacks (box size 480 pixels for 1×) were generated using resample.exe from the FREALIGN package (v9.11)^78^.

Particle alignment, 3D classification, and refinement were performed in FREALIGN. Reference maps for 30S, 50S, and 70S were generated in EMAN2^79^ by editing the 70S structure (PDB 6WDE)^80^ and low-pass filtering to 20 Å. The 4× binned stacks were first aligned to the reference with C1 symmetry over the 300–30 Å range using five global-search (mode 3) cycles, followed by five local-refinement (mode 1) cycles over the 300–18 Å range. The 2× binned stacks were then refined in mode 1 with stepwise high-resolution limits of 18, 12, 10, and 8 Å (five cycles each). The top 60% of particles by alignment score were retained to compute the 3D reconstructions. The final 2× alignment parameters were subsequently used for 3D classification of the 30S, 50S, and 70S particle sets from ΔrimM and wild-type samples. The 2× binned 30S stacks (Δ*rimM* and *wt*) were first classified into 12 classes over 50 cycles (300–8 Å resolution range). For ΔRimM, one high-resolution class containing IF1 and IF3 but lacking uS3, uS10, and uS14 (30S_Δ_*_rimM_* - state II) was taken to 1× refinement. Three additional classes were merged (87,412 particles) and underwent an additional round of 3D classification into four subclasses, yielding two high-resolution subclasses. The first subclass contained IF1 and IF3 and was missing uS3, uS10, uS13, uS14, and uS19 (30S_Δ_*_rimM_* - state I); and was selected for 1× refinement. The second subclass contained IF1 and IF3 with a complete r-proteins set representing the mature 30S subunit (30S_Δ_*_rimM_* - state III); and was selected for 1× refinement.

From the 12 wild-type classes, two were merged (137,111 particles) and subjected to focused mask (IF3 region, 30 Å radius) 3D classification into three subclasses. One high-resolution 30S subclass containing IF1 and IF3 and representing the mature 30S subunit (30S*_wt_*) was selected for 1× refinement. Additional focused classifications using spherical or 3D masks placed on the head domain or the mRNA exit tunnel did not yield further classes or improve the overall resolution.

The 2× binned 50S stacks (Δ*rimM* and *wt*) were first classified into 16 classes over 50 cycles (300–8 Å resolution range). For Δ*rimM*, eight classes lacking helix H68 (pre-50S; 223,342 particles) were merged and reclassified with a 25 Å focused mask on the RsfS region, yielding three subclasses. One high-resolution subclass with RsfS bound and helix H68 absent (pre-50S_Δ_*_rimM_* +RsfS −H68) was taken to 1× refinement. In parallel, five classes with helix H68 present (mature 50S; 255,050 particles) were merged and processed identically, producing one high-resolution subclass with RsfS and helix H68 present (50S_Δ_*_rimM_* +RsfS +H68) for 1× refinement.

Wild-type 50S data were processed in a similar way. Nine pre-50S classes lacking helix H68 were merged (625,331 particles) and reclassified with a 20 Å focused mask on the RsfS region into three subclasses. One pre-50S subclass containing partial occupancy of RsfS was picked (147,001 particles) and subjected to a second round of focused 3D classification (RsfS region, 20 Å) yielding a high-resolution subclass with RsfS and lacking helix H68 (pre-50S*_wt_* +RsfS-H68) for 1× refinement. In the same manner, three classes (152,063 particles) representing mature 50S with helix H68 present were picked and subjected to a focused mask 3D classification (RsfS region, 20 Å radius) and reclassified into three subclasses. One 50S subclass with partial occupancy of RsfS and with helix H68 present was picked (25,949 particles) and subjected to a second round of focused mask 3D classification (RsfS region, 20 Å radius). One high-resolution subclass containing RsfS and helix H68 (50S*_wt_* +RsfS +H68) was selected for 1× refinement.

The 2× binned 70S stacks (Δ*rimM* and *wt*) were first classified into 16 classes over 50 cycles (300–8 Å resolution range). For Δ*rimM*, three classes were picked, merged (223,385 particles) and reclassified with a 30 Å focused mask encompassing the P and E site tRNA regions into five subclasses. Three high-resolution subclasses were selected and merged into a single class (70S_Δ_*_rimM_*) for 1× refinement.

For wild type 70S, 11 classes were picked, merged (582,794 particles) and reclassified with a 20 Å focused mask encompassing the A and P site tRNA regions into four subclasses. One high-resolution class was picked (164,836 particles) and subjected to a second round of 3D classification using a 45 Å focused mask encompassing the P and E site tRNA regions into four subclasses. Three high-resolution subclasses were selected and merged into a single class (70S*_wt_*) for 1× refinement. However, no tRNA density was detected in any dataset, likely due to the purification with a high-salt sucrose cushion. Focused 3D classifications with a spherical mask over the tRNA sites yielded no additional classes. Likewise, masking the 30S head domain in 70S Δ*rimM* and *wt* stacks did not reveal immature head states or improve overall resolution.

The particles assigned to final high-resolution classes of 30S, 50S and 70S Δ*rimM* and *wt* were extracted from the 2x binned stack (with > 50% occupancy and > 0 score) using merge_classes.exe (part of FREALIGN distribution)^78^. Final refinements of the corresponding unbinned (1×) particle stacks were carried out using mode 1 over 5 cycles. For each, 95% of particles with the highest scores were used to generate the final 3D reconstructions. This process yielded high-resolution cryo-EM maps at ∼3.0 Å for 30S_Δ_*_rimM_* - state I (12,314 particles); at ∼2.9 Å for 30S_Δ_*_rimM_* - state II (39,565 particles); at ∼2.9 Å for 30S_Δ_*_rimM_* - state III (30,144 particles); at ∼3.1 Å for 30S*_wt_* (43,157 particles); at ∼2.7 Å for pre-50S_Δ_*_rimM_* +RsfS −H68 (52,567 particles); at ∼2.7 Å for 50S_Δ_*_rimM_* +RsfS +H68 (51,537 particles); at ∼2.7 Å for pre-50S*_wt_* +RsfS-H68 (60,038 particles); at ∼2.9 Å for 50S*_wt_* +RsfS +H68 (11,115 particles); at ∼2.6 Å for 70S_Δ_*_rimM_* (101,932 particles) and at ∼3.0 Å for 70S*_wt_* (146,167 particles), according to the 0.143 Fourier Shell Correlation (FSC) criterion, as detailed in Supplementary Tables 3 and 4, and Supplementary Figs. 5, 13. These final cryo-EM maps were used for model building and structure refinements. Local-resolution filtering was applied to the resulting cryo-EM maps, using blocres and blocfilt from the Bsoft (vs. 1.9.1) package^81^. The sharpening of the resulting cryo-EM maps was performed with bfactor.exe using a B-factor of −30 to −50 Å^2^ to interpret high-resolution details. FSC curves were calculated with FREALIGN for even and odd particle half-sets.

### Model building and refinement

The starting models for refining the 30S, 50S, and 70S ribosomes were derived from the 70S cryo-EM structure (PDB 6WDE)^80^, which served as the template. The models of IF1, IF3 and RsfS were generated using AlphaFold^82^ based on *E. coli* sequences using template structures from PDB 5LMV^43^ and PDB 6SJ6^26^, respectively. The structural models were initially fitted into the respective cryo-EM maps using rigid-body fitting in Chimera^83^, while the manual modelling was performed using Coot^84^. Poorly defined regions within the cryo-EM maps such as rRNA segments, r-proteins or domain regions, were omitted from the models when necessary. All structures were refined using phenix.real_space_refine in Phenix^85^. Secondary-structure and base-pairing restraints were applied to rRNA, when necessary. Correlation coefficients (model-to-map fit in Phenix) were also manually checked to avoid over-fitting of the models to the cryo-EM maps. FSC values between the final models and cryo-EM maps were computed in Phenix using phenix.mtriage, showing strong agreement between the structural models and cryo-EM maps.

The final models exhibit good stereochemistry, with minimal deviations in bond lengths and angles (Supplementary Tables 3, 4). Model quality was evaluated with MolProbity^86^. Structural superpositions and figures were prepared in ChimeraX^87^ and PyMOL (v2.3.1, Schrödinger).

## Data availability

The EM density maps generated in this study have been deposited in the EMDB under the following accession codes: EMD-55171 (30S_Δ_*_rimM_* - State I); EMD-55173 (30S_Δ_*_rimM_* - State II); EMD-55174 (30S_Δ_*_rimM_* – State III); EMD-55176 (pre-50S_Δ_*_rimM_*); EMD-55177 (50S_Δ_*_rimM_*); EMD-55178 (70S_Δ_*_rimM_*); EMD-55181 (30S*_wt_*); EMD-55182 (pre-50S*_wt_*); EMD-55183 (50S*_wt_*); and EMD-55185 (70S*_wt_*). The atomic coordinates generated in this study have been deposited in the PDB under the accession codes 9SS0 (30S_Δ_*_rimM_* - State I); 9SS1 (30S_Δ_*_rimM_* - State II); 9SS2 (30S_Δ_*_rimM_* – State III); 9SS4 (pre-50S_Δ_*_rimM_*); 9SS5 (50S_Δ_*_rimM_*); and 9SS6 (70S_Δ_*_rimM_*). Mass spectrometry proteomics data were deposited to the ProteomeXchange Consortium via the PRIDE partner repository under dataset identifier PXD068732.

## Acknowledgements

We wish to express our gratitude to Dr. Sylva Brabencova and Veronika Konopova for their assistance with the purification of ribosomal samples. We also appreciate the valuable comments and feedback on the manuscript from members of our laboratories. CIISB, Instruct-CZ Centre of the Instruct-ERIC EU consortium, funded by MEYS CR infrastructure project LM2023042 and the European Regional Development Fund-Project “Innovation of Czech Infrastructure for Integrative Structural Biology” (No. CZ.02.01.01/00/23_015/0008175), is gratefully acknowledged for the financial support of the measurements at the CEITEC Proteomics Core Facility. We gratefully acknowledge the Cryo-electron microscopy and tomography core facility CEITEC MU of CIISB, Instruct-CZ Centre, supported by MEYS CR (LM2023042) and the European Regional Development Fund-Project Innovation of Czech Infrastructure for Integrative Structural Biology” (No. CZ.02.01.01/00/23_015/0008175). Computational resources were provided by the e-INFRA CZ project (ID:90254), supported by MEYS CR. This work was supported from the National Institute of Virology and Bacteriology (Program EXCELES, ID Project No. LX22NPO5103), funded by the European Union - Next Generation EU (to G.D.) and by the US NIH grants GM134931, AI139202, and AG082005 (to Y.M.H.).

## Author Contributions

Conceptualization: Y.M.H. and G.D. Methodology: A.H.H., Y.N., H.G., I.M., M.P., S.N., J.D., G.B., Y.M.H. and G.D. Validation: A.H.H., Y.N., H.G., I.M., Y.M.M., and G.D. Investigation: A.H.H., Y.N., H.G., I.M., M.P., G.B., Y.M.H. and G.D. Resources: G.B., Y.M.H. and G.D. Writing - Original Draft: A.H.H., Y.N., Y.M.H. and G.D. Writing - Review and Editing: All; Visualization: A.H.H., Y.N., Y.M.H. and G.D. Supervision: Y.M.H. and G.D. Funding acquisition: Y.M.H. and G.D.

## Conflict of Interest Disclosure

The authors declare no conflict of interest.

## Notes

### Competing Interest Statement

The authors have declared no competing interest.

