## Supplementary material for "Mechanistic insights into *E. coli* recovery from growth arrest"

**Supplementary Table 1. Primers used for strain constructions**

| Gene | Primer | Sequence (5′ -> 3′) | Purpose |
|---|---|---|---|
| *rsfS* | rsfS-F | ACCCAGGGGGAAAACTTGCAGGGTAAAGCACGTGTAGGCTGGAGCTGCTTC | Δ*rsfS* strain construction |
| | rsfS-R | TTCACGCATTAACTCCAGAGTTTTTCCAGTATGGGAATTAGCCATGGTCC | |
| | rsfS-F | GGCGACAATTGTCCCGAAATCG | *rsfS* deletion check |
| | rsfS-R | GGAATTTCAATCAGCTCGAAGG | |
| *rsmE* | rsmE-F | CGACGCGGATTTTTAACTATGCGTATCCCCCGCATTTATCGTGTAGGCTGGAGCTGCTTC | Δ*rsmE* strain construction |
| | rsmE-R | CTTCTCCGTTAGCCCAAATCGCCAAATCGTACTTGTAGCGATGGGAATTAGCCATGGTCC | |
| | rsmE-F | GCAGCTGTTCAACGCATGGAAC | *rsmE* deletion check |
| | rsmE-R | CACCATTGATCAGATACAGATCG | |

**Supplementary Table 2. Primers used for plasmid constructions**

Mutated nucleotides are underlined.

| Gene | Primer | Sequence (5′ -> 3′) | Purpose |
|---|---|---|---|
| *rimM* | rimM-F | GCGTGGAAAGATCTGATGGG | Y106A/Y107A in His-*rimM* |
| | rimM-R | AGCATCACCTTCTTCCAGCTGTG | |
| *rbfA* | rbfA-F | AAAGGGCCATGGCGAAAGAATTTGGTCG | *rbfA* cloning |
| | rbfA-R | AAAGGGGGATCCTTAGTCCTCCTTGCTGTCGTCC | |

**Supplementary Table 3**

Refinement statistics for cryo-EM structures $30S_{\Delta rimM}$ - state I, $30S_{\Delta rimM}$ state - II, $30S_{\Delta rimM}$ state - III, pre-$50S_{\Delta rimM}$ +RsfS (-H68), $50S_{\Delta rimM}$ +RsfS +H68 and $70S_{\Delta rimM}$

| | **$30S_{\Delta rimM}$ class I** | **$30S_{\Delta rimM}$ class II** | **$30S_{\Delta rimM}$ class III** | **pre-$50S_{\Delta rimM}$ +RsfS -H68** | **$50S_{\Delta rimM}$ +RsfS +H68** | **$70S_{\Delta rimM}$** |
|---|---|---|---|---|---|---|
| **PDBID** | 9SS0 | 9SS1 | 9SS2 | 9SS4 | 9SS5 | 9SS6 |
| **EMBD** | EMD-55171 | EMD-55173 | EMD-55174 | EMD-55176 | EMD-55177 | EMD-55178 |
| **Data collection and processing** | | | | | | |
| Magnification | 105,000x | 105,000x | 105,000x | 105,000x | 105,000x | 105,000x |
| Voltage (kV) | 300 | 300 | 300 | 300 | 300 | 300 |
| Electron exposure (e–/Å2) | 40 | 40 | 40 | 40 | 40 | 40 |
| Defocus range (µm) | -0.5 to -2.0 | -0.5 to -2.0 | -0.5 to -2.0 | -0.5 to -2.0 | -0.5 to -2.0 | -0.5 to -2.0 |
| Pixel size (Å) | 0.834 | 0.834 | 0.834 | 0.834 | 0.834 | 0.834 |
| Symmetry imposed | C1 | C1 | C1 | C1 | C1 | C1 |
| Initial particle (no.) | 455076 | 455076 | 455076 | 652894 | 652894 | 900281 |
| Final particle (no.) | 12314 | 39565 | 30144 | 52567 | 51537 | 101932 |
| Map resolution (Å) | 3.0 | 2.9 | 2.9 | 2.7 | 2.7 | 2.6 |
| FSC threshold | 0.143 | 0.143 | 0.143 | 0.143 | 0.143 | 0.143 |
| **Refinement** | | | | | | |
| Initial model used (PDB code) | 6WDE | 6WDE | 6WDE | 6WDE | 6WDE | 6WDE |
| Model resolution (Å) | 3.0 | 3.0 | 3.0 | 3.0 | 3.0 | 3.0 |
| Correlation Coefficient (cc_mask)* | 0.84 | 0.89 | 0.88 | 0.91 | 0.92 | 0.90 |
| Map sharpening B factor ($Å^2$) | -30 | -30 | -30 | -40 | -40 | -50 |
| Model composition* | | | | | | |
| Non-hydrogen atoms | 47157 | 49977 | 53770 | 91562 | 92721 | 144897 |
| Protein residues | 1968 | 2190 | 2630 | 3698 | 3584 | 6015 |
| RNA residues | 1475 | 1525 | 1539 | 2928 | 3023 | 4562 |
| B factors ($Å^2$)* | | | | | | |
| Protein | 157.09 | 182.47 | 182.00 | 150.16 | 127.89 | 138.97 |
| RNA | 170.19 | 183.44 | 146.71 | 135.18 | 126.31 | 124.61 |
| R.m.s. deviations*,§ | | | | | | |
| Bond lengths (Å) | 0.004 | 0.004 | 0.004 | 0.004 | 0.004 | 0.004 |
| Bond angles (°) | 0.769 | 0.716 | 0.758 | 0.702 | 0.700 | 0.717 |
| Validation# | | | | | | |
| MolProbity score | 1.94 | 1.83 | 1.86 | 1.47 | 1.49 | 1.61 |
| Clashscore | 6.26 | 5.01 | 5.45 | 3.06 | 3.10 | 3.36 |
| Poor rotamers (%) | 0.37 | 0.16 | 0.14 | 0.40 | 0.14 | 0.30 |
| Ramachandran plot# | | | | | | |
| Favored (%) | 88.21 | 89.35 | 89.44 | 94.57 | 94.18 | 92.21 |
| Allowed (%) | 11.63 | 10.51 | 10.44 | 5.37 | 5.77 | 7.69 |
| Disallowed (%) | 0.16 | 0.14 | 0.12 | 0.06 | 0.06 | 0.10 |
| Validation (RNA)# | | | | | | |
| Good sugar pucker (%) | 98.85 | 99.21 | 98.83 | 99.21 | 99.14 | 99.17 |
| Good backbone (%) | 80.14 | 82.75 | 84.86 | 84.15 | 83.96 | 85.14 |

* from Phenix
### from Molprobity
§ root mean square deviations

**Supplementary Table 4**

Refinement statistics for cryo-EM maps $30S_{wt}$, pre-$50S_{wt}$ +RsfS (-H68), $50S_{wt}$ +RsfS +H68 and $70S_{wt}$

| | **$30S_{wt}$** | **pre-$50S_{wt}$ +RsfS (-H68)** | **$50S_{wt}$ +RsfS +H68** | **$70S_{wt}$** |
|---|---|---|---|---|
| **EMBD** | EMD-55181 | EMD-55182 | EMD-55183 | EMD-55185 |
| **Data collection and processing** | | | | |
| Magnification | 105,000x | 105,000x | 105,000x | 105,000x |
| Voltage (kV) | 300 | 300 | 300 | 300 |
| Electron exposure (e–/Å2) | 40 | 40 | 40 | 40 |
| Defocus range (μm) | -0.5 to -2.0 | -0.5 to -2.0 | -0.5 to -2.0 | -0.5 to -2.0 |
| Pixel size (Å) | 0.834 | 0.834 | 0.834 | 0.834 |
| Symmetry imposed | C1 | C1 | C1 | C1 |
| Initial particle (no.) | 826581 | 970120 | 970120 | 978750 |
| Final particle (no.) | 43157 | 60038 | 11115 | 146167 |
| Map resolution (Å) | 3.1 | 2.7 | 2.9 | 3.0 |
| FSC threshold | 0.143 | 0.143 | 0.143 | 0.143 |

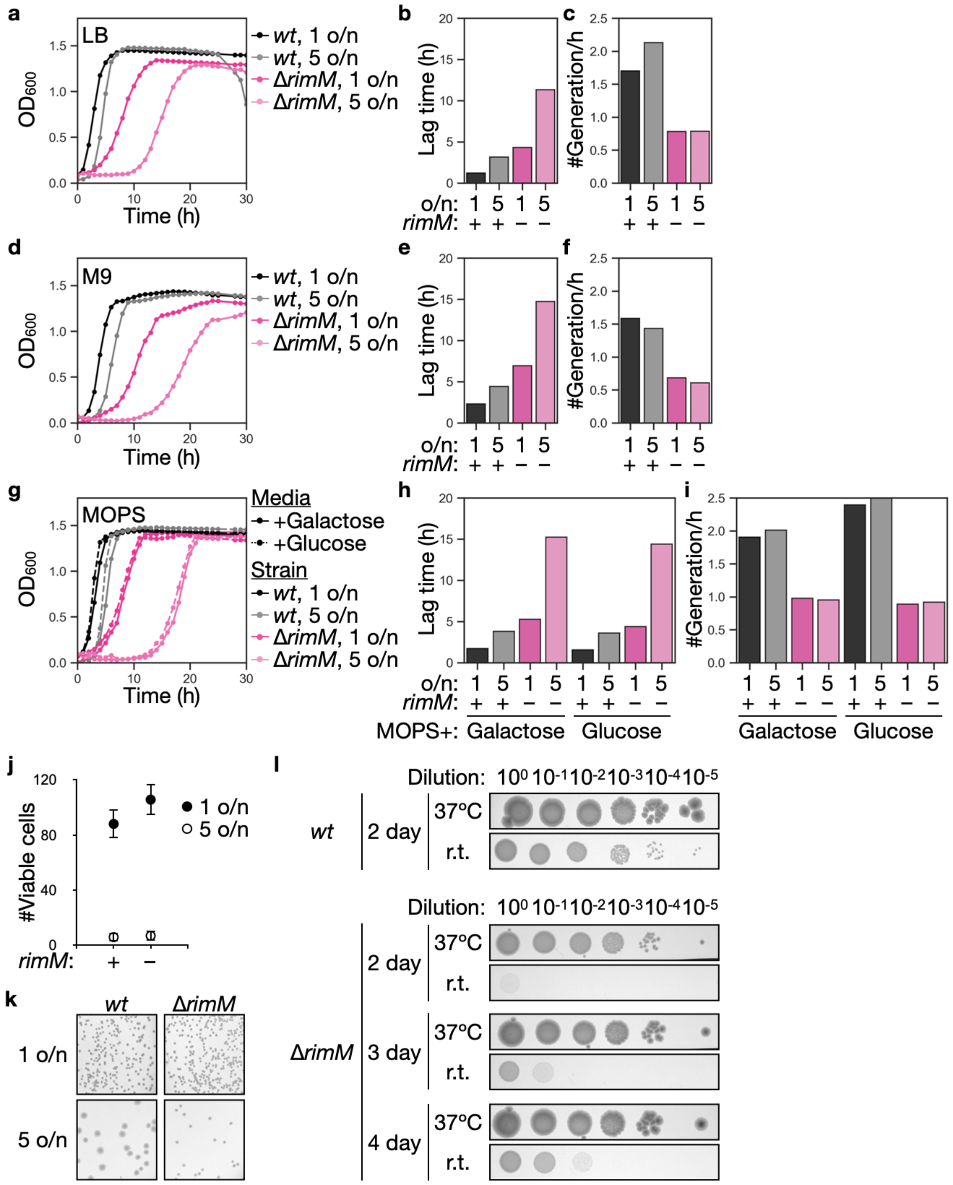

**Supplementary Figure 1. Assembly of *E. coli* 30S subunit in growth recovery. (a)** Growth profiles of the *wt* and Δ*rimM* strains of MG1655 in LB at 37ºC after 1-day or 5-day pre-culture (n = 1). **(b)** Lag time calculated from $OD_{600}$'s of cultures in (a), as the time required to reach $OD_{600}$ of 0.2. **(c)** Growth rates (the number of generations per hour) of cultures in (a). **(d)** Growth profiles of *wt* and Δ*rimM* strains of MG1655 in M9 minimal medium at 37ºC after 1-day or 5-day pre-culture (n = 1). **(e)** Lag time calculated from $OD_{600}$'s of cultures in (d). **(f)** Growth rates (the number of generations per hour) of cultures in (d). **(g)** Growth profiles of *wt* and Δ*rimM* strains of MG1655 in MOPS rich medium supplemented with 0.2% galactose or 0.2% glucose at 37ºC after 1-day or 5-day pre-culture (n = 1). **(h)** Lag time calculated from $OD_{600}$'s of cultures in (g). **(i)** Growth rates (the number of generations per hour) of cultures in (g). **(j)** Viable cells of *wt* and Δ*rimM* strains of MG1655 (n = 6; technical replicates). Cell counts were calculated from the colony number within an area of the same size on LB plates spread after continuous liquid culture for 1 day or 5 days at 37ºC. **(k)** Zoomed-in views of *wt* and Δ*rimM* strains of MG1655 on LB plates after 1-day or 5-day liquid culture. **(l)** Cold sensitivity of *wt* and Δ*rimM* strains of MG1655 after one overnight growth in LB and incubated on LB plates at room temperature (r.t.) or 37ºC.

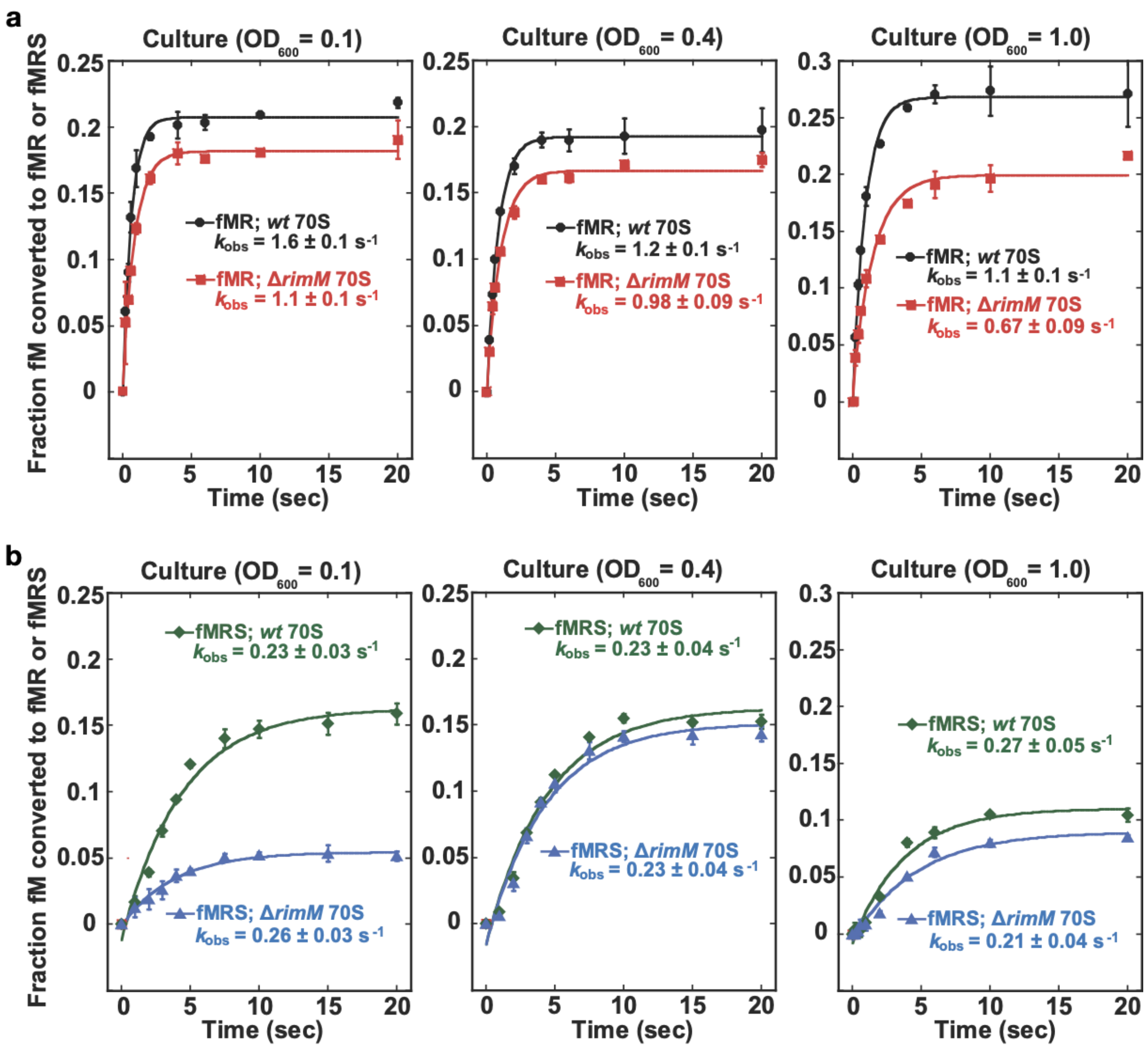

**Supplementary Figure 2. Kinetics of synthesis of dipeptide and tripeptide by *E. coli wt* and Δ*rimM* ribosomes. (a)** A time course of the fractional conversion of fM to fMR by Δ*rimM* and *wt* MRE600 ribosomes harvested at the indicated $OD_{600}$'s. Error bars, mean ± SD (n = 3). **(b)** A time course of the conversion of fM to fMRS by Δ*rimM* and *wt* ribosomes harvested at the indicated $OD_{600}$'s. Error bars, mean ± SD (n = 3).

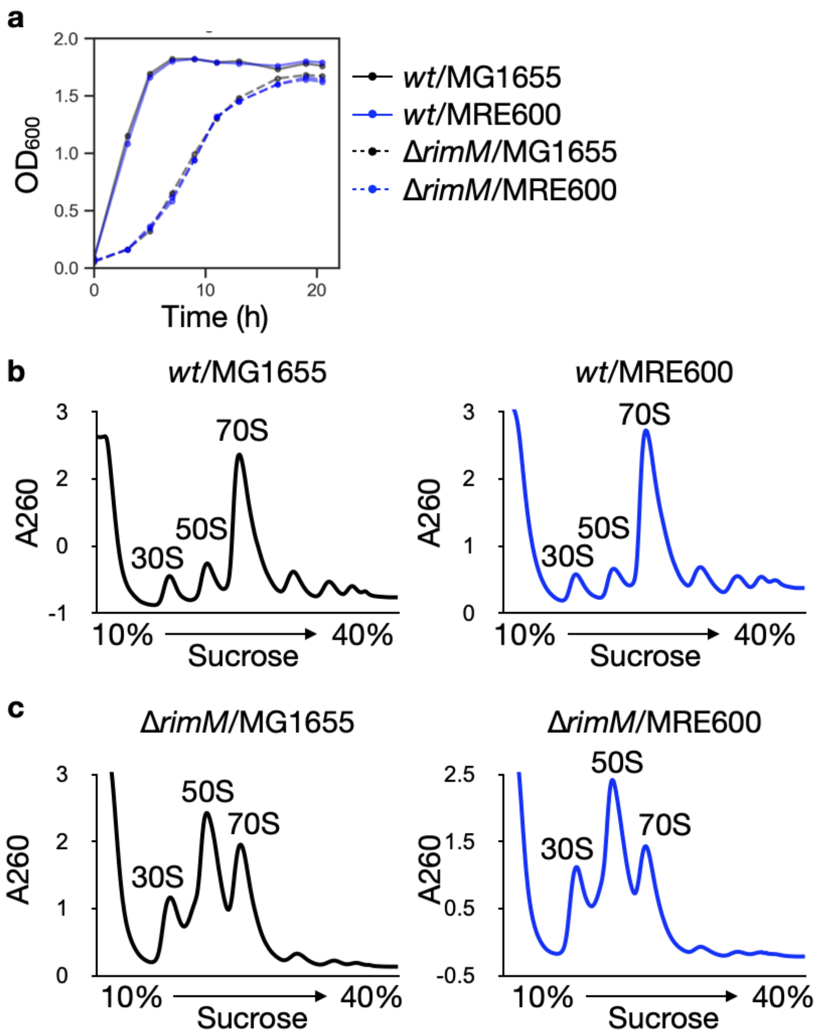

**Supplementary Figure 3. Similar profiles of *E. coli* MG1655 and MRE600. (a)** Growth profiles of *wt* and Δ*rimM* strains in MG1655 and in MRE600 in LB (n = 1). **(b)** Polysome profiles of *wt* strains in MG1655 and in MRE600 harvested at $OD_{600}$ 0.4 in LB (n = 1) and fractionated in a 10-40% sucrose gradient. **(c)** Polysome profiles of Δ*rimM* strains in MG1655 and in MRE600 harvested at $OD_{600}$ 0.4 in LB (n = 1) and fractionated in a 10-40% sucrose gradient.

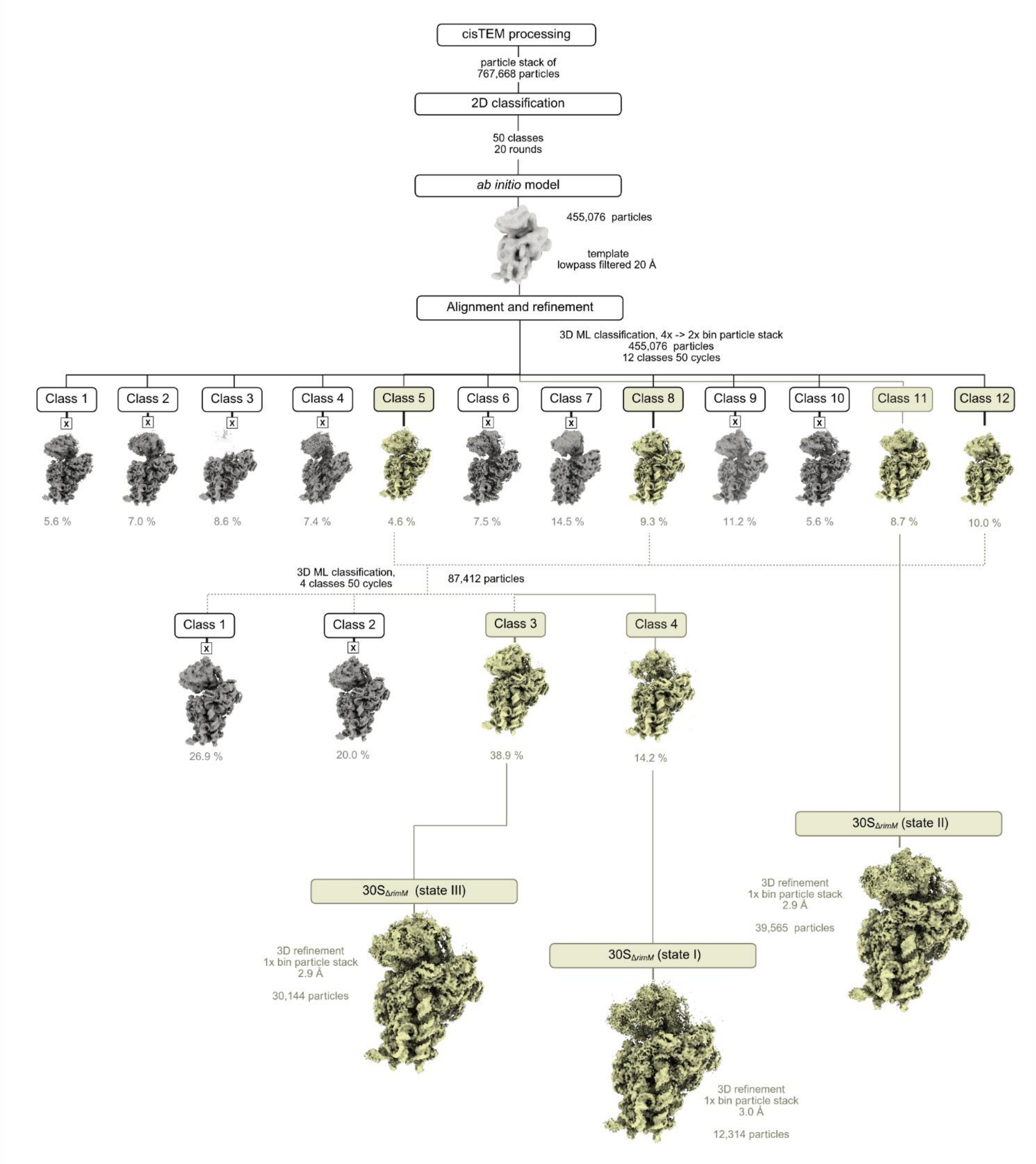

**Supplementary Figure 4. Cryo-EM data processing and classification scheme for 30S$_{\Delta rimM}$ structures.** Low-resolution and junk classes are shown in grey, while classes advancing to 3D classification are shown in yellow. Final maps used for structural modelling are displayed at the bottom, along with the number of particles and corresponding resolutions (FSC = 0.143).

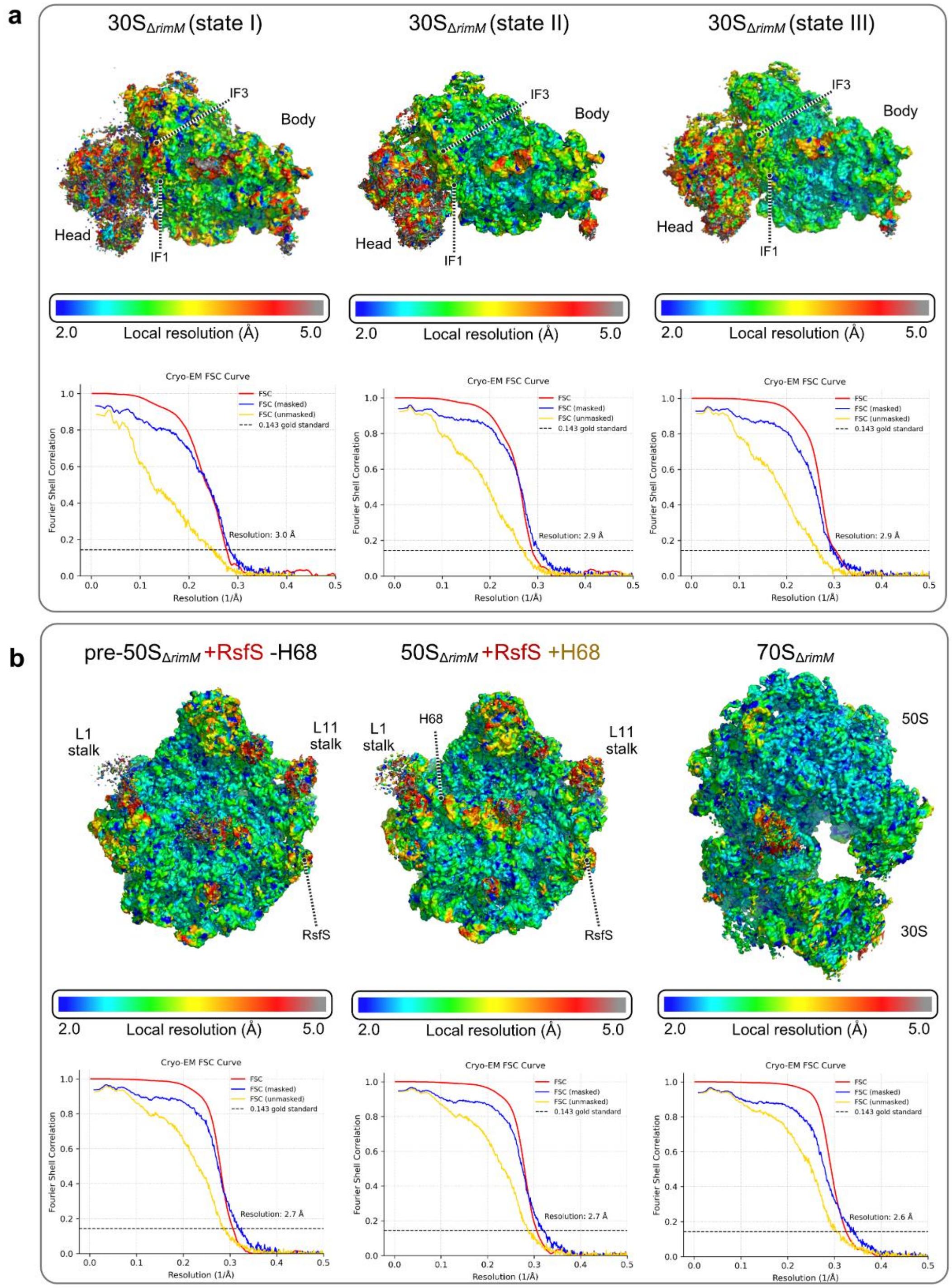

**Supplementary Figure 5. Global and local resolution for Δ*rimM* 30S structures.** (**a**) Local resolutions in respective $30S_{\Delta rimM}$ cryo-EM maps (top row). Fourier shell correlation (FSC) between even- and odd-particle half maps (red) reveals that map resolutions range from 2.9 to 3.0 Å (at FSC = 0.143, dotted line); FSC between final models and final maps (cyan) masked and unmasked FSCs are also shown (bottom row). (**b**) Local resolutions in respective $50S_{\Delta rimM}$ and $70S_{\Delta rimM}$ cryo-EM maps (top row). Fourier shell correlation (FSC) between even- and odd-particle half maps (red) reveals that map resolutions range from 2.9 to 3.0 Å (at FSC = 0.143, dotted line); FSC between final models and final maps (cyan) masked and unmasked FSCs are also shown (bottom row).

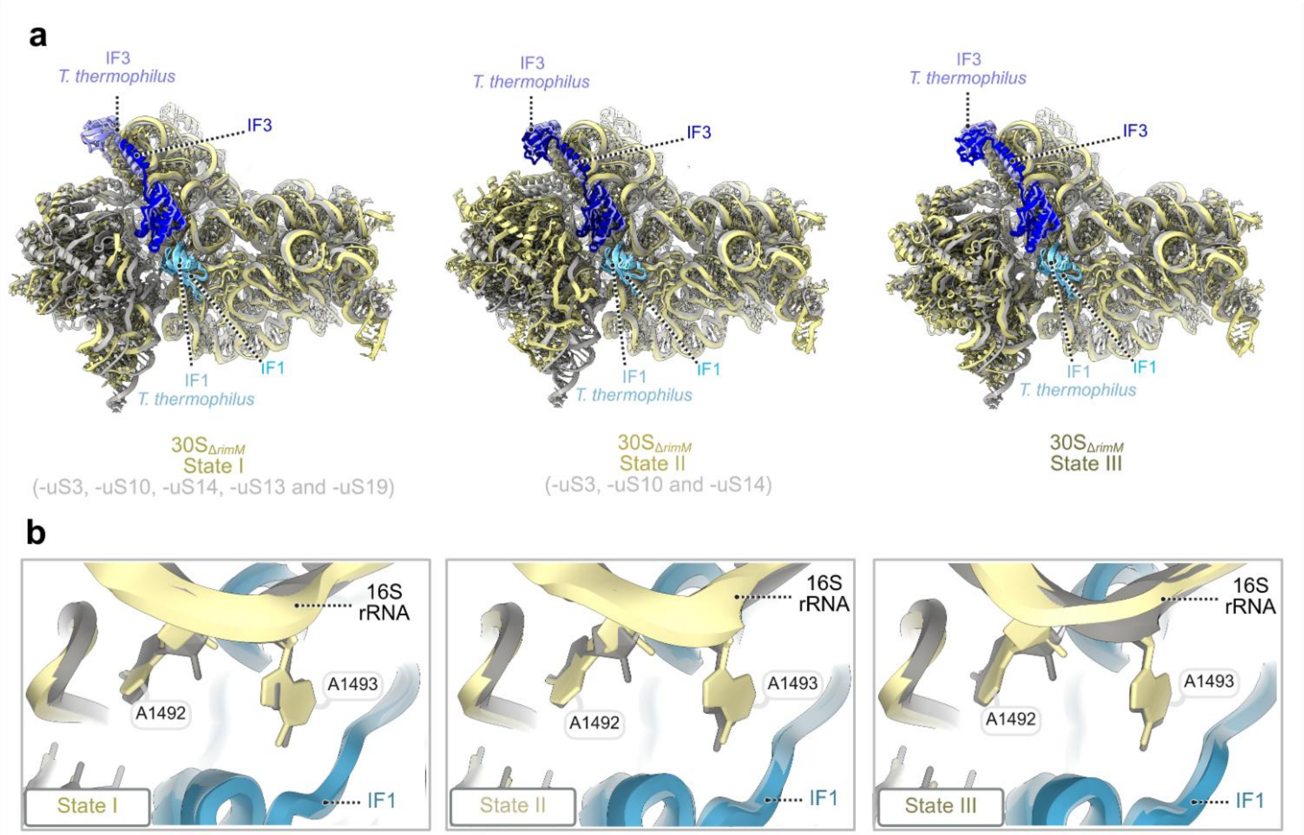

**Supplementary Figure 6. Structural comparison of 30S$_{\Delta RimM}$ states to the *T. thermophilus* 30S initiation complex. (a)** Superposition of 30S$_{\Delta rimM}$ states (I-III) with the 30S initiation complex (PDB: 5LMN) shows that IF1 and IF3 hold their canonical positions on the 30S subunit. **(b)** Close-up view of the decoding site illustrates that the arrangement of rRNA helices and decoding nucleotides A1492 and A1493 in Δ*rimM* states matches that of the initiation complex (PDB: 5LMN). Structural models of 30S$_{\Delta rimM}$ subunit (state I -III) are shown in yellow, while initiation complex of *T. thermophilus* in grey (PDB: 5LMN). The initiation factors IF1 (light blue for state I, II and III and cyan for *T. thermophilus*) and IF3 (dark blue for state I, II and III and light purple for *T. thermophilus*) are also highlighted. Structural alignments were performed based on the 16S rRNA localized in the 30S body for each pairwise comparison.

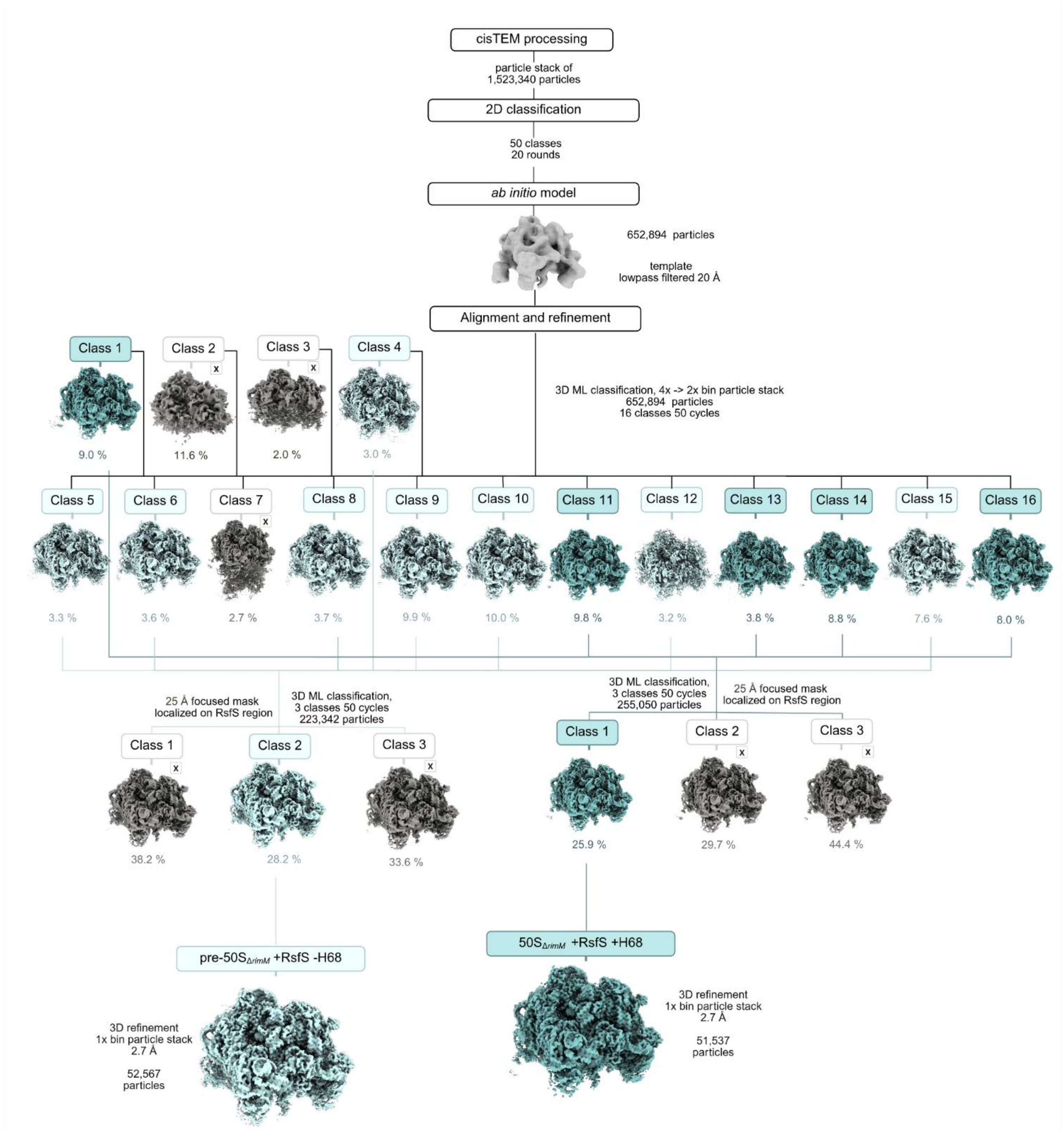

**Supplementary Figure 7. Cryo-EM data processing and classification scheme for 50S$_{\Delta rimM}$ structures.** Low-resolution and junk classes are shown in grey, while classes advancing to 3D classification are shown in light and dark cyan. Final maps used for structural modelling are displayed at the bottom, along with the number of particles and corresponding resolutions (FSC = 0.143).

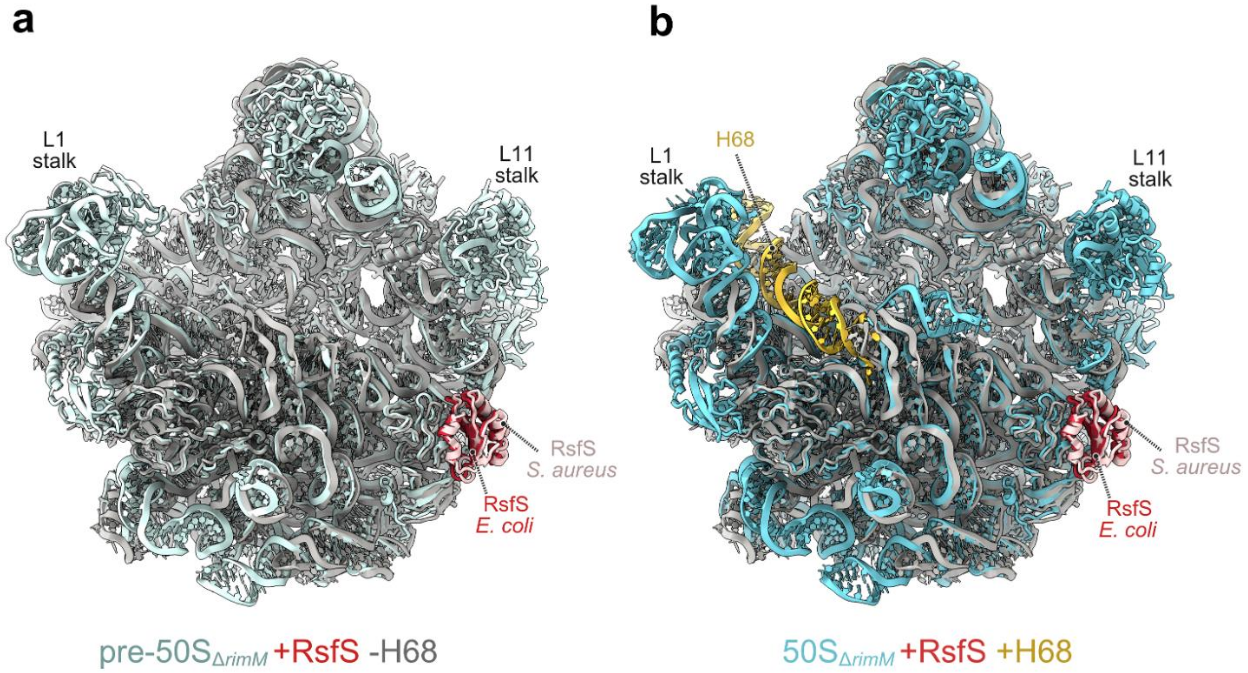

**Supplementary Figure 8. Structural comparison of $50S_{\Delta RimM}$ states to the *S. aureus* 50S-RsfS complex. (a)** Superposition of pre-$50S_{\Delta rimM}$ state with the *S. aureus* 50S–RsfS complex (PDB: 6SJ6) shows consistent positioning of RsfS in the structures. **(b)** Superposition of $50S_{\Delta rimM}$ state with the *S. aureus* 50S–RsfS complex (PDB: 6SJ6) shows consistent positioning of RsfS in the structures. The pre-$50S_{\Delta rimM}$ ribosomal subunit is shown in light blue, while $50S_{\Delta rimM}$ is in cyan, aligned to 50S–RsfS complex from *S. aureus* in grey (PDB: 6SJ6). All structures are shown in crown view from the intersubunit interface, with RsfS highlighted in red for the $50S_{\Delta rimM}$ states and light pink for *S. aureus* RsfS. H68 in $50S_{\Delta rimM}$ is highlighted in gold. Structural alignments were performed based on the 23S rRNA core of the 50S subunit for each pairwise comparison.

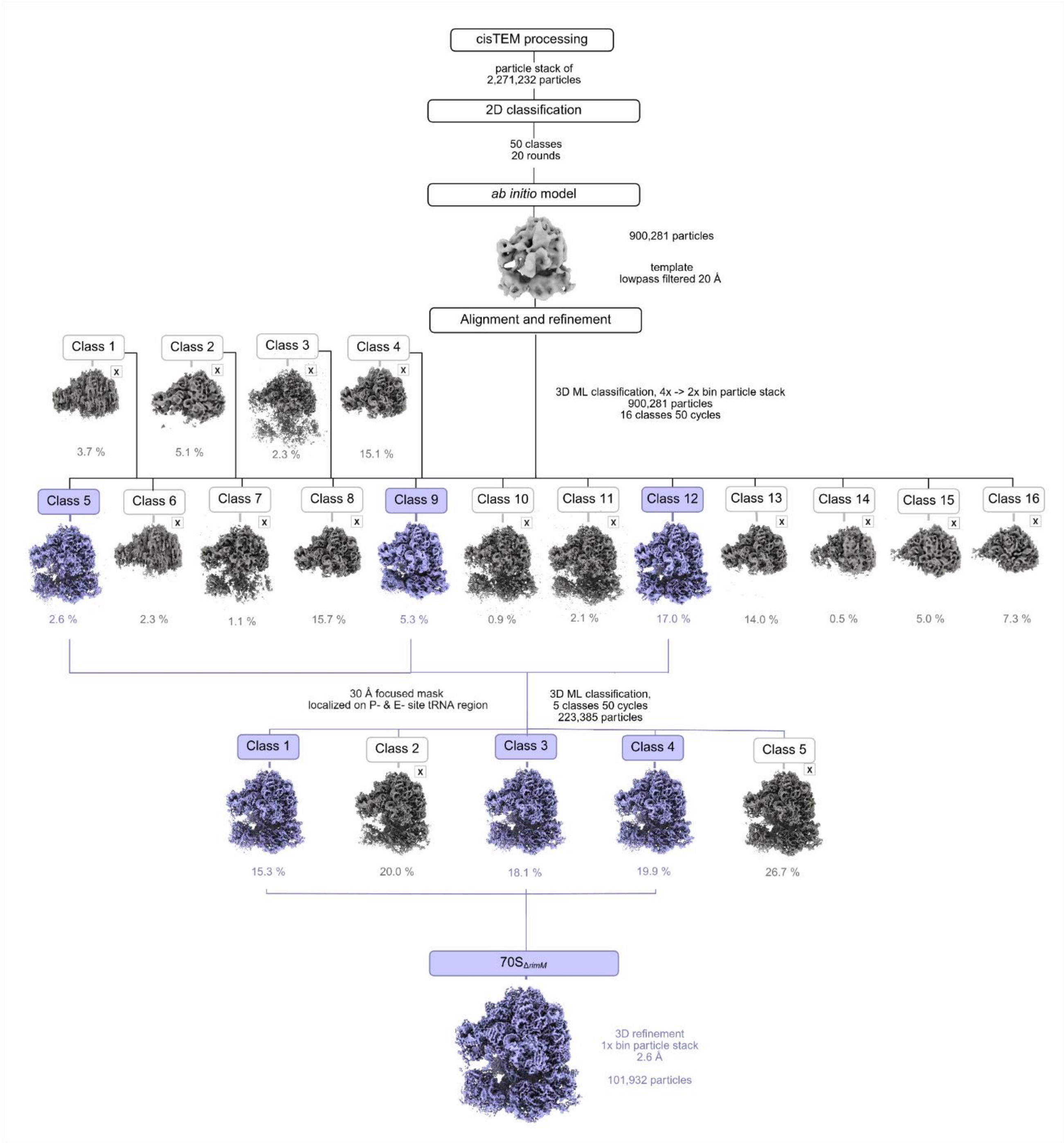

**Supplementary Figure 9. Cryo-EM data processing and classification scheme for 70S$_{\Delta rimM}$ structure.** Low-resolution and junk classes are shown in grey, while classes advancing to 3D classification are shown in light and dark cyan. Final maps used for structural modelling are displayed at the bottom, along with the number of particles and corresponding resolutions (FSC = 0.143).

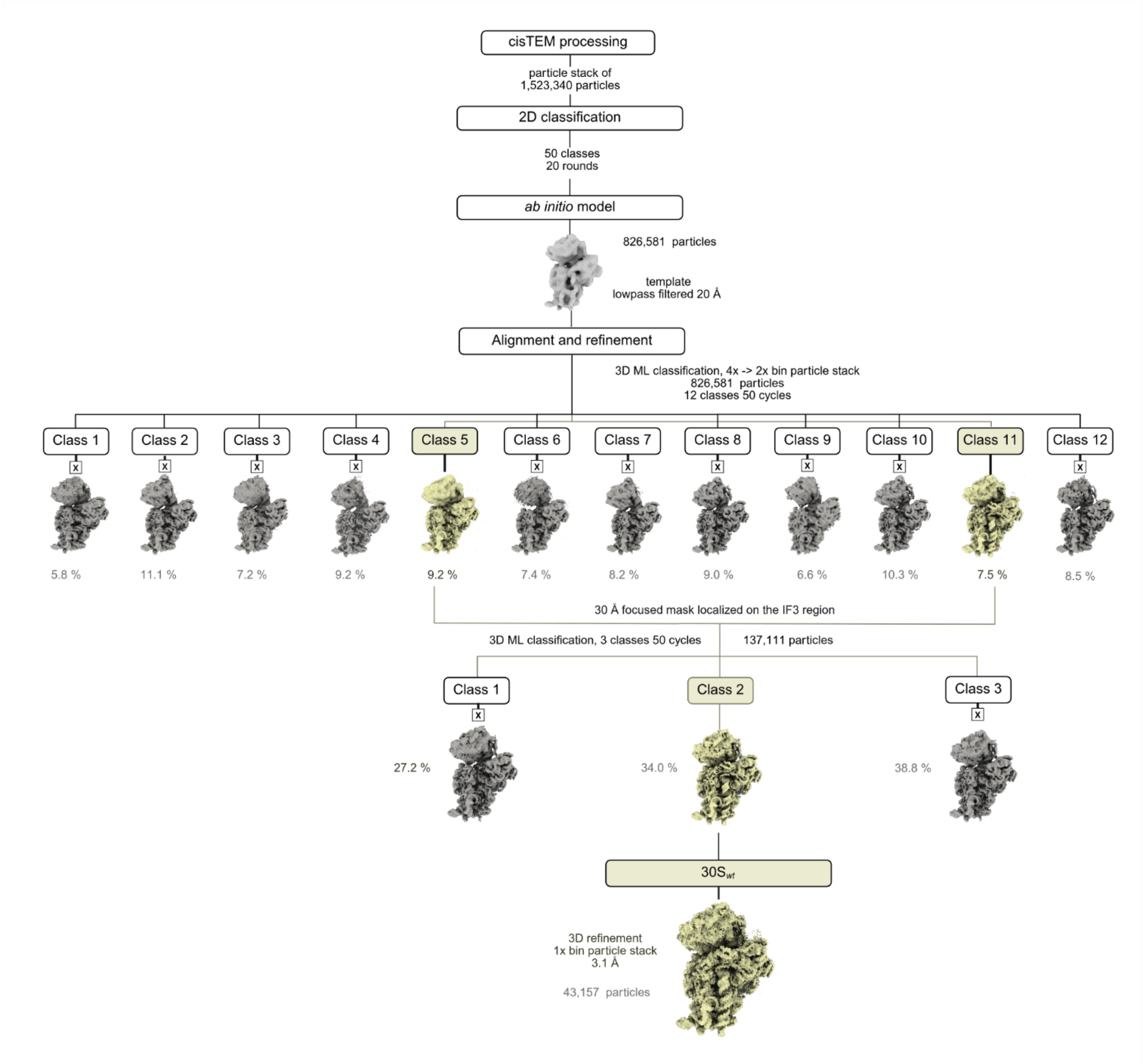

**Supplementary Figure 10. Cryo-EM data processing and classification scheme for 30S$_{wt}$ structure.** Low-resolution and junk classes are shown in grey, while classes advancing to 3D classification are shown in yellow. Final maps used for structural modelling are displayed at the bottom, along with the number of particles and corresponding resolutions (FSC = 0.143).

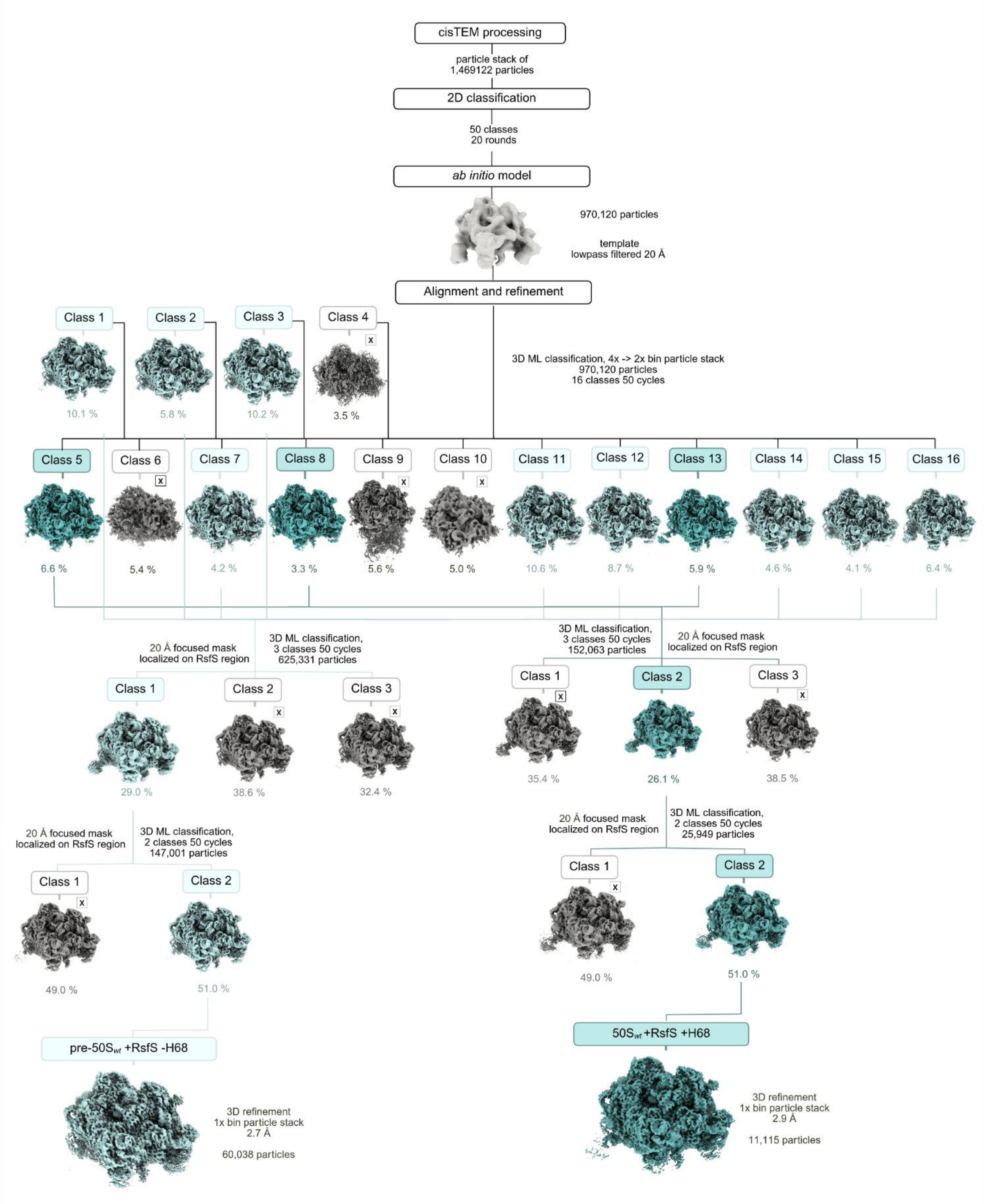

**Supplementary Figure 11. Cryo-EM data processing and classification scheme for 50S$_{wt}$ structures.** Low-resolution and junk classes are shown in grey, while classes advancing to 3D classification are shown in light and dark cyan. Final maps used for structural modelling are displayed at the bottom, along with the number of particles and corresponding resolutions (FSC = 0.143).

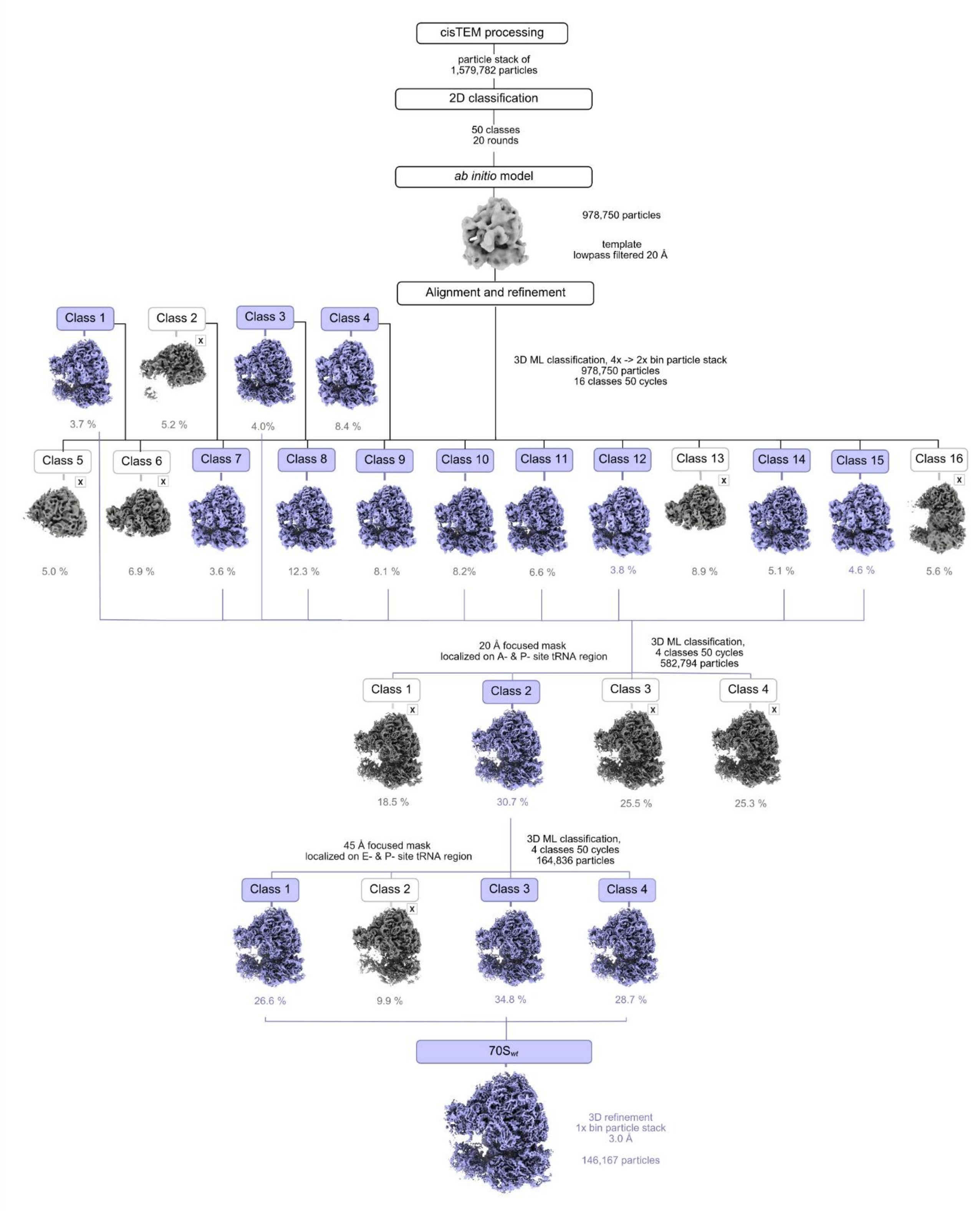

**Supplementary Figure 12. Cryo-EM data processing and classification scheme for 70S*wt* structure.** Low-resolution and junk classes are shown in grey, while classes advancing to 3D classification are shown in light and dark cyan. Final maps used for structural modelling are displayed at the bottom, along with the number of particles and corresponding resolutions (FSC = 0.143).

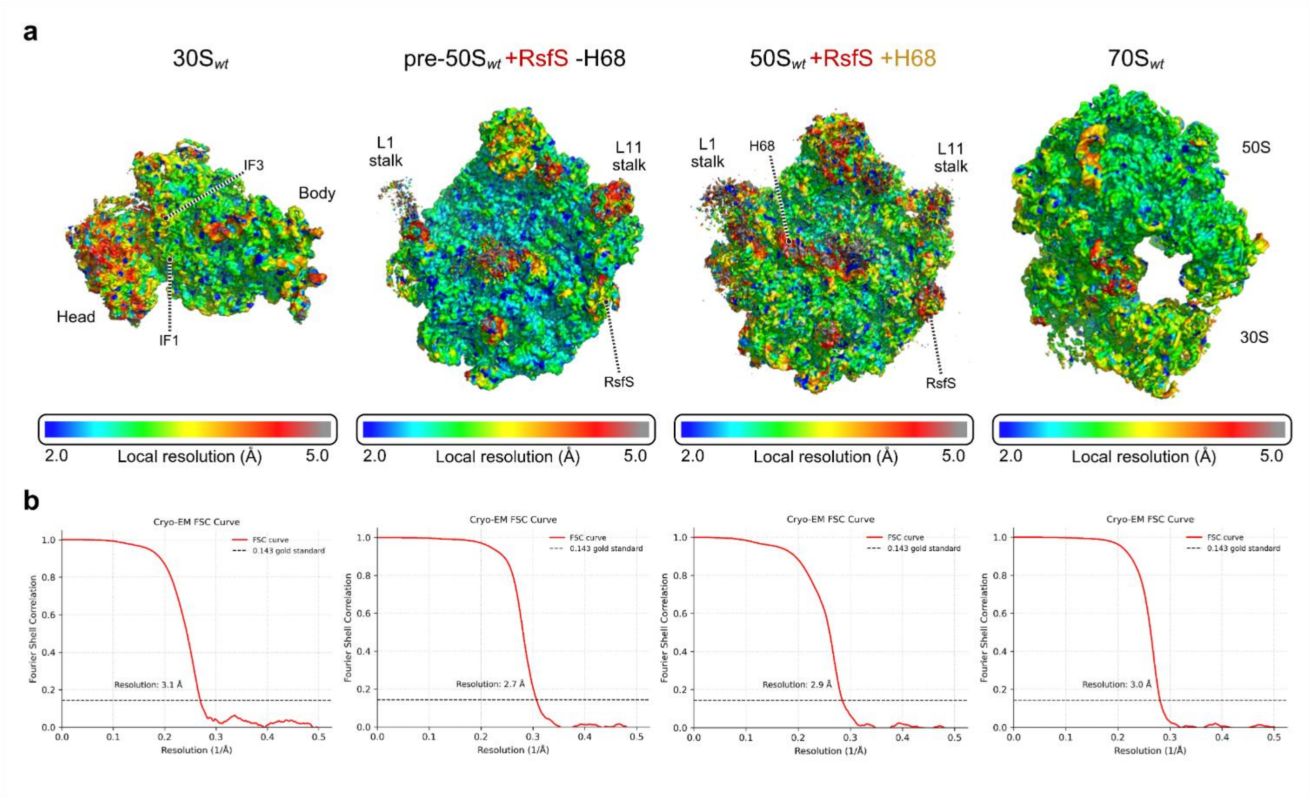

**Supplementary Figure 13. Global and local resolution for wild type structures. (a)** Local resolutions in respective cryo-EM maps. **(b)** Fourier shell correlation (FSC) between even- and odd-particle half maps (red) shows that map resolutions range from 2.9 to 3.0 Å (at FSC = 0.143, dotted line); FSC between final models and final maps (cyan) masked and unmasked FSCs are also shown.

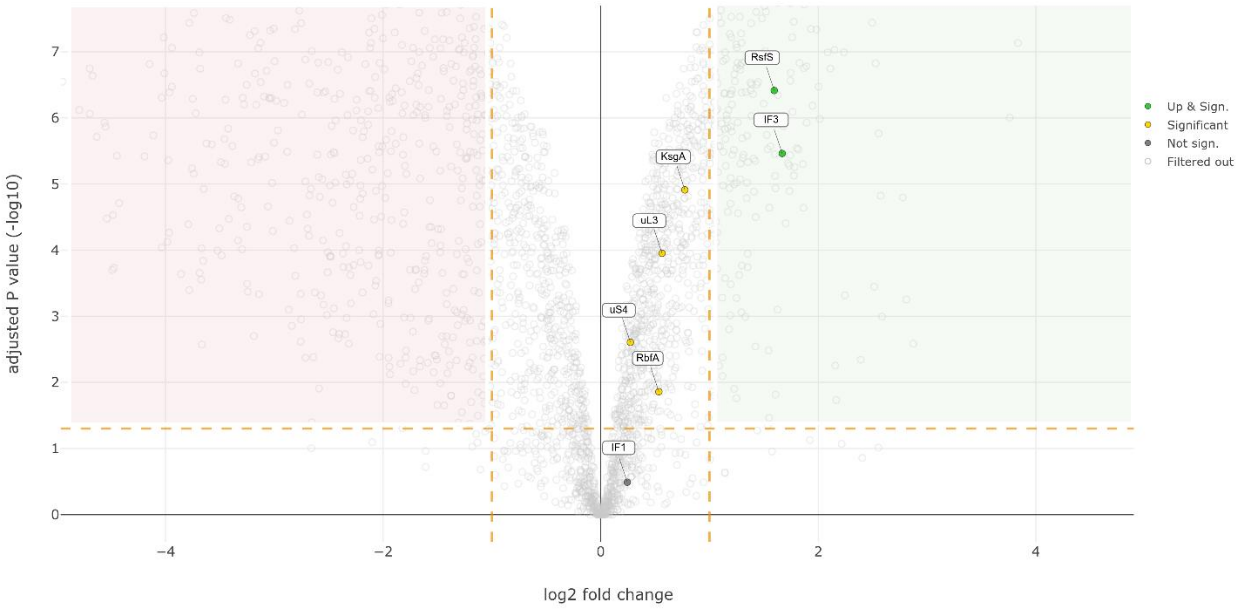

**Supplementary Figure 14. Differential protein expression analysis of *wt* and Δ*rimM* strains by mass spectrometry.** Volcano plot showing the log2 fold change in protein abundance (x-axis) versus the adjusted p-value (−log10, y-axis) for the comparison between *wt* and Δ*rimM* MRE600 strains at $OD_{600}$ of 0.4. Proteins highlighted in green segment are significantly upregulated in Δ*rimM* (adjusted p-value < 0.05, log2 fold change > 1), while yellow dots indicate proteins that are statistically significant but do not meet the fold-change cutoff. Grey dots represent non-significant proteins, and light grey open circles indicate filtered-out data points. Key ribosome-associated proteins, including RsfS and IF3, are highlighted, showing strong upregulation, while ribosomal proteins uS4, uL3 and 30S maturation factors RbfA and KsgA display modest increases, and IF1 shows no upregulation. The dashed lines mark the thresholds for significance.

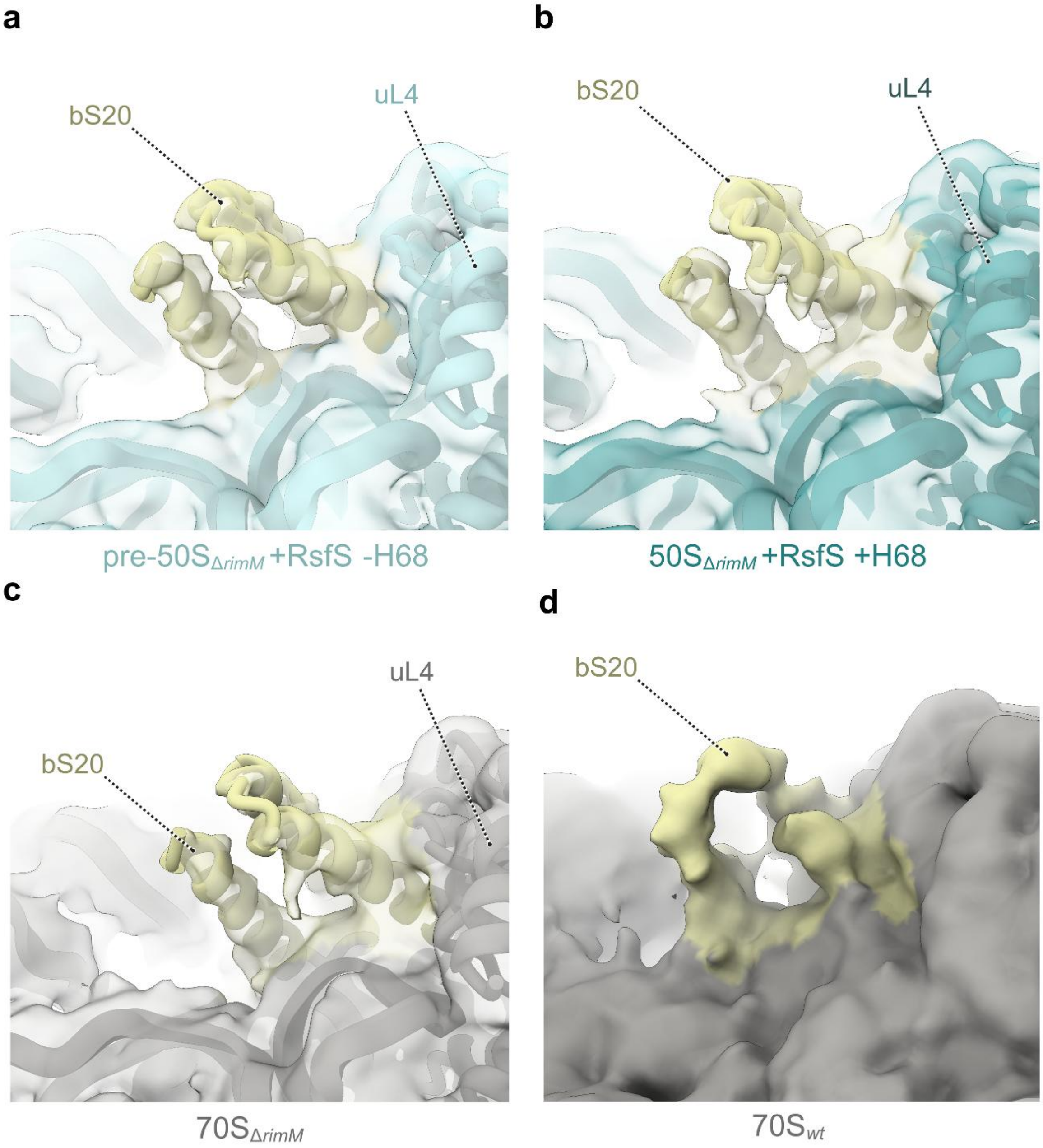

**Supplementary Figure 15. Visualization of bS20 in 50S subunits and 70S ribosomes. (a)** Close-up view on pre-50S$_{\Delta rimM}$ subunit (light cyan) highlighting the location of bS20 (yellow). The cryo-EM map (transparent surface) is shown at 1.5 σ. **(b)** Close-up view of the 50S$_{\Delta rimM}$ subunit (cyan) highlighting the location of bS20 (yellow). The cryo-EM map is shown at 1.5 σ. **(c)** Close-up view on the 70S$_{\Delta rimM}$ in the uL4 region of the 50S subunit (grey) highlighting the location of bS20 (yellow). The cryo-EM map is shown at 1.0 σ. **(d)** Close-up view on the 70S$_{wt}$ cryo-EM map (grey surface) highlighting the location of bS20 (yellow surface). The cryo-EM map is shown at 1.0 σ. In all cases, the cryo-EM maps were low-pass filtered applying a B-factor of 100 Å$^2$.
